# *Chlamydomonas* RABL4/IFT27 Mediates Phototaxis via Promoting BBSome-dependent Ciliary Export of Phospholipase D

**DOI:** 10.1101/2022.02.06.479262

**Authors:** Yan-Xia Liu, Bin Xue, Zhen-Chuan Fan

## Abstract

Phospholipase D (PLD) interacts with the BBSome for loading onto retrograde intraflagellar transport (IFT) trains to exit cilia in *Chlamydomonas reinhardtii*. PLD ciliary retention and depletion of Rab-like 4 (RABL4) GTPase IFT27 cause the same non-phototactic phenotype but not impair IFT and ciliation. Here, we show that the IFT-B1 subunit IFT27 binds its partner IFT25 to form the heterodimeric IFT25/27 in an IFT27 nucleotide state-independent manner. IFT25/27, IFT-A, and IFT-B are irrelevant for maintaining the stability of one another. GTP-loading onto IFT27 enhances the IFT25/27 affinity for IFT-B1 in cytoplasm, while GDP-loaded IFT27 does not prevent IFT25/27 from entering and cycling through cilia by integrating into IFT-B1. Upon at the ciliary tip, IFT25/27 cycles on and off IFT-B1 and this process is irrelevant with the nucleotide state of IFT27. During BBSome remodeling at the ciliary tip, IFT25/27 promotes BBSome reassembly independent of IFT27 nucleotide state, making post-remodeled BBSomes available for PLD to interact with. Therefore, IFT25/27 facilitates BBSome-dependent PLD export from cilia via controlling availability of intact BBSomes at the ciliary tip, providing a regulatory mechanism for IFT27 to mediate phototaxis in *C. reinhardtii*.

## Introduction

The cilium consists of microtubule-assembled axoneme covered by a specialized ciliary membrane (Rosenbaum & Witman, 2002). As a signaling antenna of eukaryotic cells, the cilium protrudes from the cell surface and senses and transmits extracellular stimuli inside the cell. Underlying the ciliary sensory role is the fact that many G protein-coupled receptors (GPCRs), ion channels, and their downstream signaling proteins specifically distribute to the ciliary membrane (Wingfield *et al*, 2018). These signaling proteins cycle between ciliary membrane and cell membrane through motor protein-driven intraflagellar transport (IFT) trains along the axoneme (Kozminski *et al*, 1993; Wingfield *et al*., 2018). IFT trains consist of 22 different IFT proteins organized into IFT-A (6 subunits) and IFT-B (16 subunits) sub-complexes (Cole *et al*, 1998; Follit *et al*, 2009). IFT-B is further divided into biochemically and functionally different core IFT-B1 and peripheral IFT-B2 entities (Lucker *et al*, 2005; Taschner *et al*, 2014; Taschner *et al*, 2016). In a dynamic state, kinesin-2 motor binds IFT-B1 to drive anterograde IFT trains from the ciliary base toward the ciliary tip (anterograde IFT) (Cole *et al*., 1998). Instead, cytoplasmic dynein 1b (cytoplasmic dynein 2 in mammals) returns retrograde IFT trains in an opposite direction via coupling with IFT-A (retrograde IFT) (Pazour *et al*, 1998). During IFT, the BBSome composed of eight Bardet-Biedl syndrome (BBS) proteins (BBS1/2/4/5/7/8/9/18) (seven proteins including BBS1/2/4/5/7/8/9 in *Chlamydomonas reinhardtii*) links specific signaling proteins as cargoes to IFT trains for maintaining their ciliary dynamics (Lechtreck *et al*, 2009a; Nachury *et al*, 2007; Wingfield *et al*., 2018).

As an IFT cargo adaptor, the BBSome delivers signaling proteins for either entering or exiting cilia via IFT (Wingfield *et al*., 2018). BBSome malfunction thus causes loss or abnormal retention of signaling proteins in the ciliary membrane (Chiang *et al*, 2004; Lechtreck *et al*., 2009a; Loktev *et al*, 2008; Nachury *et al*., 2007; Scheidecker *et al*, 2014; Sun *et al*, 2021; Xue *et al*, 2020; Zhang *et al*, 2011). These defects impair signaling protein dynamics in cilia and eventually leads to Bardet-Biedl syndrome (BBS) in humans (Fliegauf *et al*, 2007) and phototactic defects in *C. reinhardtii* (Lechtreck *et al*., 2009a; Liu & Lechtreck, 2018). Our previous study has identified the BBSome disassembles first at the ciliary tip followed by reassembling into intact entities when turning around the ciliary tip during IFT (Sun *et al*., 2021). During this remodeling process, signaling proteins interact with the post-remodeled BBSome for loading onto retrograde IFT trains followed by exporting from cilia via IFT. Thus far, small GTPases of various subfamilies have been found to play roles in regulating BBSome remodeling and BBSome-cargo interaction in cilia. For instance, *Chlamydomonas* Rab-like 4 (RABL4) GTPase IFT27 enables BBSome reassembly at the ciliary tip and Arf-like 6 (ARL6) GTPase BBS3 instead promotes BBSome interaction with phospholipase D (PLD) for its export form cilia (Liu *et al*, 2021b; Sun *et al*., 2021). As with different mechanisms for controlling BBSome availability for signaling protein loading for their export form cilia via IFT, IFT27 and BBS3 dysfunction both can disrupt BBSome ciliary signaling, eventually causing phototactic defects in *C. reinhardtii* (Liu *et al*., 2021b; Sun *et al*., 2021).

Unlike other conventional IFT-B1 subunits essential for IFT-B assembly, IFT initiation, and ciliation (Fan *et al*, 2010; Hou *et al*, 2007), IFT25 and IFT27, by binding to form a heterodimer IFT25/27 (Wang *et al*, 2009), are dispensable for IFT-B assembly nor for ciliation in rodent somatic cells and green algae (Dong *et al*, 2017b; Eguether *et al*, 2014; Keady *et al*, 2012; Liew *et al*, 2014; Sun *et al*., 2021). In rodent cells, knockout of IFT27 or IFT25 causes abnormal ciliary retention of BBSomes as well as signaling proteins like patched-1 (PTCH1), smoothened (Smo), Gli2, and GPR161 due to the defect in BBSome ciliary removal (Eguether *et al*., 2014; Keady *et al*., 2012; Liew *et al*., 2014). As expected, siblings carrying IFT27 mutants displayed BBS due to hedgehog signaling dysfunction (Aldahmesh *et al*, 2014; Schaefer *et al*, 2019). Underlying this observation, IFT25/27 detaches from IFT-B at the ciliary tip and binds and activates BBS3 as a BBS3-specific guanine nucleotide exchange factor (GEF) (Liew *et al*., 2014). IFT27 was thus proposed to promote BBSome loading onto retrograde IFT trains via the BBS3 pathway (Jin *et al*, 2010; Liew *et al*., 2014). A second model proposed IFT25/27 to do so by providing the BBSome a docking site in IFT-B of retrograde IFT trains (Eguether *et al*., 2014). In contrast, our previous study has shown that *Chlamydomonas* IFT25/27 promotes BBSome reassembly at the ciliary tip rather than directly enables BBSome loading onto retrograde IFT trains or provides the BBSome a docking site in IFT-B1, suggesting that *Chlamydomonas* IFT27 differs from its rodent counterpart in maintaining BBSome dynamics in cilia (Liu *et al*., 2021b; Sun *et al*., 2021).

PLD relies on IFT/BBS for moving out of cilia and BBSome dysfunction impairs PLD dynamics in cilia, causing phototactic defects in *C. reinhardtii* (Liu & Lechtreck, 2018). IFT27 deficiency disrupts BBSome reassembly at the ciliary tip and leads to the non-phototactic phenotype (Sun *et al*., 2021), while whether IFT27 mediates phototaxis via PLD remains to be established. Besides, whether and how nucleotide state of IFT27 plays a role in regulating phototactic behavior remains elusive. By performing functional, *in vivo* biochemical, and single-particle *in vivo* imaging assays, we showed that IFT27 mediates phototaxis via promoting BBSome-dependent ciliary removal of PLD but in a nucleotide-independent manner in *C. reinhardtii*.

## Results

### The stability of IFT-A, IFT-B, and IFT25/27 are irrelevant

IFT-B is composed of the core IFT-B1 and the peripheral IFT-B2 subcomplexes (Lucker *et al*., 2005; Taschner *et al*., 2014; Taschner *et al*., 2016). Our study and others have shown that the IFT-B stability relies on the proper integration of several of its IFT-B1 core proteins (Brazelton *et al*, 2001; Fan *et al*., 2010; Hou *et al*., 2007; Pazour *et al*, 2000), while the IFT-B1 core proteins IFT25 and IFT27 form a heterodimer (IFT25/27) which occasionally dissociates from IFT-B1 but not influences the stability of IFT-B (Dong *et al*., 2017b; Sun *et al*., 2021; Wang *et al*., 2009). To have a full view on the interplay among IFT-A, IFT-B, and IFT25/27, we monitored IFT-A subunits IFT43 and IFT139, IFT-B1 subunits IFT46 and IFT70, IFT-B2 subunits IFT38 and IFT57 as well as IFT25 and IFT27 in the IFT-B1 subunit null mutants *bld1*/*ift52* and *bld2/ift46-1* (Brazelton *et al*., 2001; Hou *et al*., 2007). In *bld1*/*ift52*, significantly reduced abundances of IFT38, IFT46, IFT57, and IFT70 were observed yet with unchanged amounts of IFT43, IFT139, IFT25, and IFT27 (***Figure 1A, B***) (Brazelton *et al*., 2001). In *bld2*/*ift46-1*, IFT46 was nondetectable as expected, while the amounts of IFT38, IFT57, and IFT70 were reduced significantly with unchanged abundances of IFT43, IFT139, IFT25, and IFT27 (***Figure 1A, B***) (Hou *et al*., 2007). In agreement with these observations at protein level, both mutants had significantly elevated abundances of IFT43 and IFT139 mRNAs along with unchanged abundances of IFT38, IFT46 (only for *bld1/ift52*), IFT57, IFT70, IFT25, and IFT27 transcripts (***Figure 1C***). These results suggest that depletion of a *bona fida* IFT-B1 subunit causes the dissociation and instability of IFT-B complex but not affects the stability of IFT-A and IFT25/27 in *C. reinhardtii* (Hou *et al*., 2007; Pazour *et al*., 2000). We next performed the reciprocal experiment in the IFT-A null mutants *ift43* and *ift139* (Zhu *et al*, 2017). Knockout of IFT43 did not alter the abundances of IFT38, IFT46, IFT57, IFT70, IFT25, and IFT27 both at protein and transcript levels but caused significantly reduced abundance of IFT139 yet with elevated IFT139 transcript (***Figure 1D-F***). The same pattern was also observed in *ift139* mutant (***Figure 1D-F***). Thus, much like what was observed for IFT-B, the stability of IFT-A relies on the proper integration of several of its component proteins (Zhu *et al*., 2017). As IFT-B and IFT25/27 complex proteins remain unchanged during IFT-A depletion, the stability of IFT-B and IFT25/27 does not depend on the presence of IFT-A. Taken together, the biochemical stability of IFT-A and IFT-B are independent of each other and do not affect IFT25/27 stability in *C. reinhardtii*. As reflected by our previous studies that depletion of IFT25 or IFT27 alone does not alter the abundance of IFT-A and IFT-B (Dong *et al*., 2017b; Sun *et al*., 2021). Given that knockdown of IFT25 or IFT27 alone did not alter the mRNA abundances of IFT43, IFT139, IFT46, IFT70, IFT38, and IFT57, we therefore conclude that the stability of IFT-A, IFT-B, and IFT25/27 are irrelevant in *C. reinhardtii* (***Figure 1G, H***).

**Figure 1.**
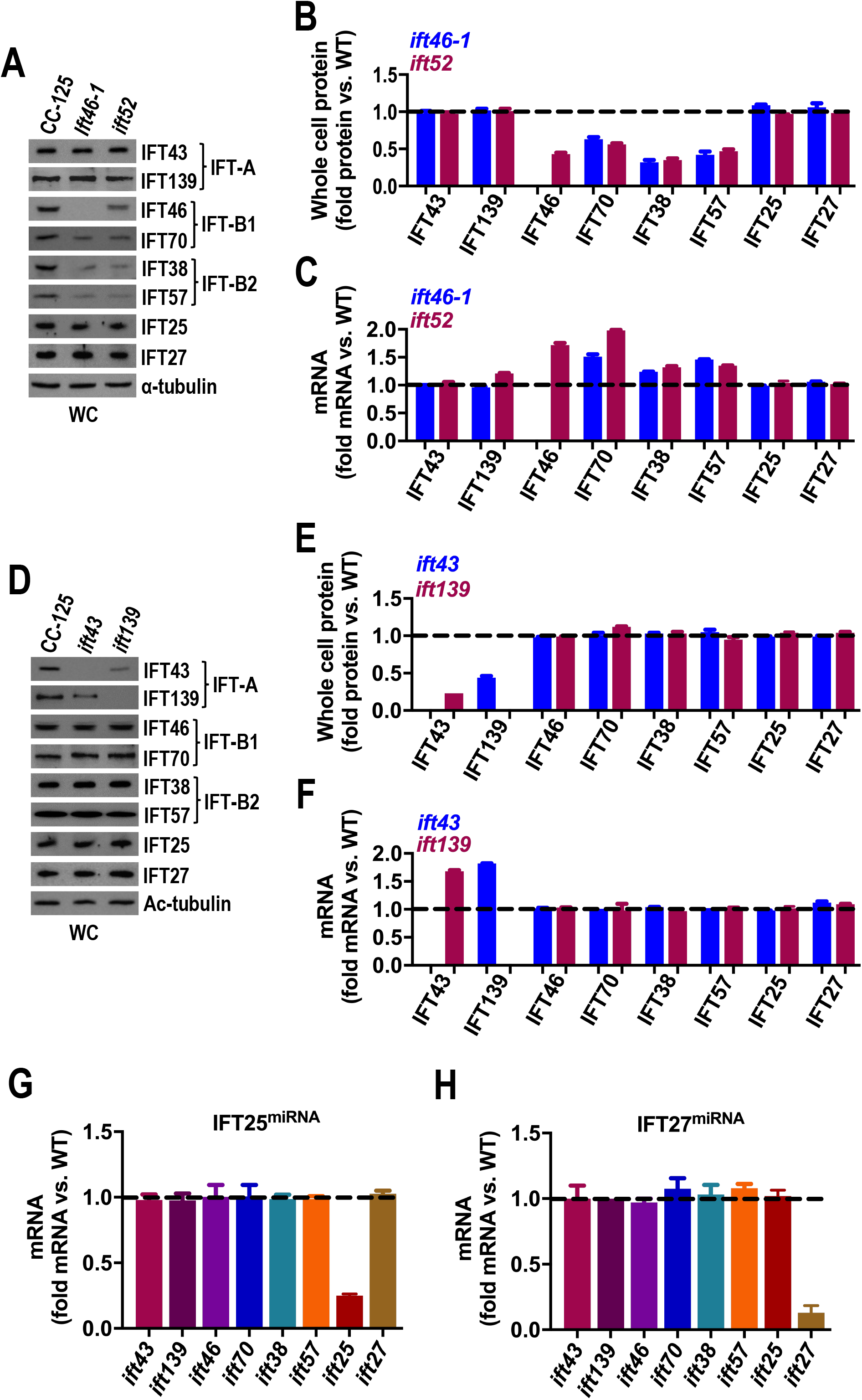
The stability of IFT-A, IFT-B, and IFT25/27 are irrelevant. (**A** and **B**) Immunoblots of whole cell (WC) samples of CC-125, *ift52*, and *ift46-1* cells probed with antibodies against the IFT proteins as shown (**A**) and quantification of the target proteins (**B**). Alpha-tubulin was used to adjust the loading. (**C**) Quantification of IFT protein mRNA levels in *ift52* and *ift46-1* cells. Messenger RNA level was normalized to the GBLP housekeeping gene and presented as fold-change relative to CC-125 mRNA. (**D** and **E**) Immunoblots of WC samples of CC-125, *ift43*, and *ift139* cells probed with antibodies against the IFT proteins as shown (**D**) and quantification of the target proteins (**E**). Alpha-tubulin was used to adjust the loading. (**F**) Quantification of IFT protein mRNA levels in *ift43* and *ift139* cells. Messenger RNA level was normalized to the GBLP housekeeping gene and presented as fold-change relative to CC-125 mRNA. (**G** and **H**) Quantification of IFT protein mRNA levels in IFT25^miRNA^ and IFT27^miRNA^ cells. Messenger RNA level was normalized to the GBLP housekeeping gene and presented as fold-change relative to CC-125 mRNA. For panels B and E, protein level was presented as fold-change relative to CC-125 protein. For panels other than A and D, each value represents the mean ± S.D. from three independent measurements of the assay.

### IFT27 is dispensable for maintaining normal IFT

IFT25 and IFT27 bind to form a heterodimer (IFT25/27), providing us the biochemical evidence that two proteins could be functionally overlapped (Dong *et al*., 2017b; Wang *et al*., 2009). Indeed, IFT25- and IFT27-null mouse show similar BBS-like phenotypes and two proteins are indispensable for ciliary hedgehog signaling (Aldahmesh *et al*., 2014; Eguether *et al*., 2014; Keady *et al*., 2012; Liew *et al*., 2014). In *Chlamydomonas* and rodent cells, IFT27 stability relies on IFT25 as an IFT27-binding partner but not *vice versa*; disruption of IFT25 thus causes IFT27 depletion which eventually deprives IFT25/27 of the ciliary presence (Dong *et al*., 2017b; Keady *et al*., 2012; Sun *et al*., 2021). Our previous studies have shown that IFT25- and IFT27-knockdown *Chlamydomonas* strains harbor normal IFT (other than IFT25/27) protein contents in cilia in a static state, resembling the IFT25- and IFT27-null rodent cells (Dong *et al*., 2017b; Eguether *et al*., 2014; Keady *et al*., 2012; Sun *et al*., 2021). To investigate whether IFT27 could affect motor protein contents in *C. reinhardtii*, we examined the IFT27-knockdown strain IFT27^miRNA^, which harbors ∼10% IFT27 protein as compared to the wild-type (WT) CC-125 strain, and the IFT27-rescuing strain IFT27^Res-WT^ with a C-terminal hemagglutinin (HA) and green fluorescence protein (GFP)-double tagged IFT27 (IFT27-HA-GFP) expressed in the IFT27^miRNA^ strain (***Figure 2A***) (Sun *et al*., 2021). The IFT27^miRNA^ and IFT27^Res-WT^ cells contained normal abundance of the motor proteins FLA10 and D1bLIC both in whole cell samples and cilia (***Figure 2A, B***). To examine the effect of IFT27 on IFT in a dynamic state, we expressed IFT43 attached at its C-terminus to HA and yellow fluorescent protein (YFP) (IFT43-HA-YFP) at similar amounts in CC-125 and IFT27^miRNA^ cells (resulting strains *IFT43 IFT43-HA-YFP* and *IFT43 IFT43-HA-YFP IFT27KD*) (***Figure 2C***). *IFT43 IFT43-HA-YFP* and *IFT43 IFT43-HA-YFP IFT27KD* cells contained similar amount of IFT43-HA-YFP in cilia (***Figure 2D***). Total internal reflection fluorescence (TIRF) imaging of living cells observed IFT43-HA-YFP to undergo similar IFT in cilia of *IFT43 IFT43-HA-YFP* and *IFT43 IFT43-HA-YFP IFT27KD* cells (***Figure 2E, F*; *Table 1; Figure 2-videos 1*** and ***2***). These data suggest that IFT27, like IFT25, is dispensable for maintaining motor protein contents nor for normal IFT in *C. reinhardtii* (Dong *et al*., 2017b).

**Table 1.**
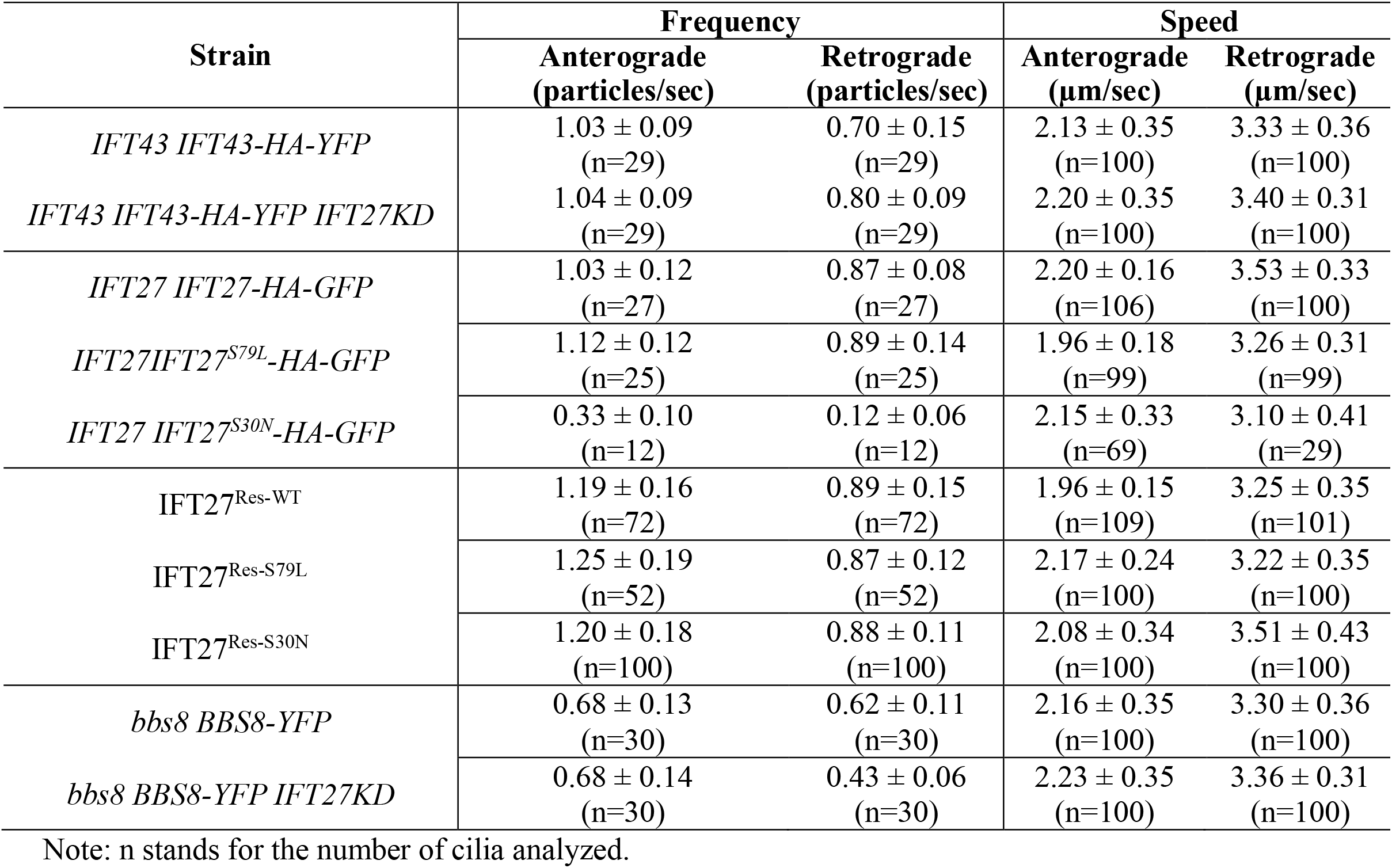
Transport of fluorescence protein-tagged particles in cilia.

**Figure 2.**
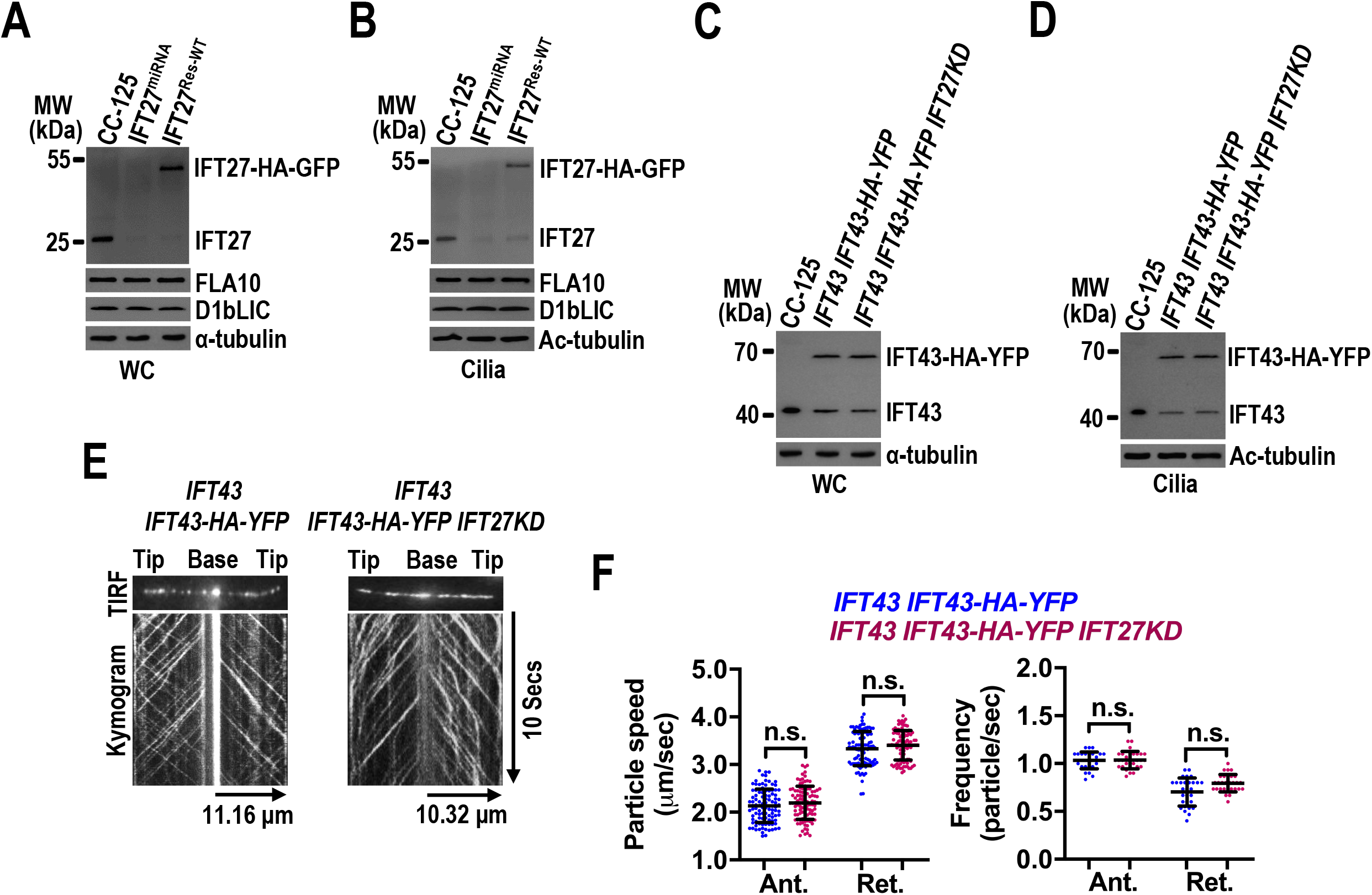
IFT27 is dispensable for maintaining normal IFT. (**A** and **B**) Immunoblots of WC samples (**A**) and cilia (**B**) of CC-125, IFT27^miRNA^ and IFT27^Res-WT^ cells with antibodies against IFT27 and the motor proteins FLA10 and D1bLIC. Knockdown of IFT27 did not alter the cellular and ciliary abundance of FLA10 and D1bLIC. (**C** and **D**) Immunoblots of WC samples (**C**) and cilia (**D**) of CC-125, *IFT43 IFT43-HA-YFP* and *IFT43 IFT43-HA-YFP IFT27KD* cells probed with α-IFT43. (**E**) TIRF images and corresponding kymograms of *IFT43 IFT43-HA-YFP* and *IFT43 IFT43-HA-YFP IFT27KD* cells (***Figure 2-videos 1*** and ***2***, 15 fps). The time and transport lengths are indicated on the right and on the bottom, respectively. (**F**) Speeds and frequencies of IFT43-HA-YFP to traffic inside cilia of *IFT43 IFT43-HA-YFP* and *IFT43 IFT43-HA-YFP IFT27KD* cells were similar. Error bar indicates S.D. n.s.: non-significance. For panels **A** and **C**, α-tubulin was used as a loading control. For panels **B** and **D**, acetylated-(Ac)-tubulin was used as a loading control.

### GTP loading only enhances IFT27’s binding affinity to IFT-B1

Although originally identified as simply a structural subunit of the IFT-B1 subcomplex, IFT27 was later reported to bind IFT-B1 as its GTP-specific effector to enter cilia in mouse and trypanosome cells (Eguether *et al*., 2014; Huet *et al*, 2014; Liew *et al*., 2014). *Chlamydomonas* IFT27 has been shown to be an active GTPase with low intrinsic GTPase activity (Bhogaraju *et al*, 2011). To “lock” IFT27 in either GDP-bound (inactive) or GTP-bound (active) state, we introduced point mutations into IFT27 that are predicted to either disrupt GTP binding while allowing limited GDP binding (IFT27^S30N^) or prevent GTP hydrolysis (IFT27^S79L^) (***Figure 3A; Figure 3-figure supplement 1A***) (Liew *et al*., 2014; Paduch *et al*, 2001; Wachter *et al*, 2019). As expected, the S79L mutation disrupted IFT27 GTP hydrolysis, suggesting that IFT27^S79L^ is a constitutive-active variant (***Figure 3-figure supplement 1B; Figure 3-source data 1***). IFT27^S30N^ did not show any GTPase activity, indicating that IFT27^S30N^ is an inactive GDP-locked or simply a dominant-negative variant (***Figure 3-figure supplement 1B; Figure 3-source data 1***), establishing that our nucleotide-binding variants are valid. To examine how the nucleotide state of IFT27 contributes to IFT-B1 binding for its ciliary entry and cycling, we expressed IFT27-HA-GFP, IFT27^S79L^-HA-GFP and IFT27^S30N^-HA-GFP at similar amount in CC-125 cells (resulting strains *IFT27 IFT27-HA GFP, IFT27 IFT27*^*S79L*^*-HA-GFP*, and *IFT27 IFT27*^*S30N*^*-HA-GFP*) (***Figure 3B***), both IFT27-HA-GFP and IFT27^S79L^-HA-GFP entered cilia, while IFT27^S30N^-HA-GFP was only detected in cilia when twenty times more ciliary extracts was loaded for immunoblotting and the blot was over-exposed after probing with anti-IFT27 (***Figure 3C***). IFT27^S79L^-HA-GFP, like IFT27-HA-GFP, immunoprecipitated IFT46 and IFT70 in cilia, while IFT27^S30N^-HA-GFP rarely recovered these proteins (***Figure 3D***). Concordant with this biochemical evidence, IFT27-HA-GFP and IFT27^S79L^-HA-GFP underwent IFT at similar speeds and frequencies (***Figure 3E, F; Table 1; Figure 3-videos 1*** and ***2***) (Dong *et al*., 2017b; Ishikawa *et al*, 2014; Qin *et al*, 2007). Only very weak signal was detected for IFT27^S30N^-HA-GFP in cilia; the faint signals indicated that it traffics at normal speeds but with strongly reduced frequencies for both anterograde and retrograde IFT as compared to IFT27-HA-GFP and IFT27^S79L^-HA-GFP (***Figure 3E, F; Table 1; Figure 3-video 3***).

**Figure 3.**
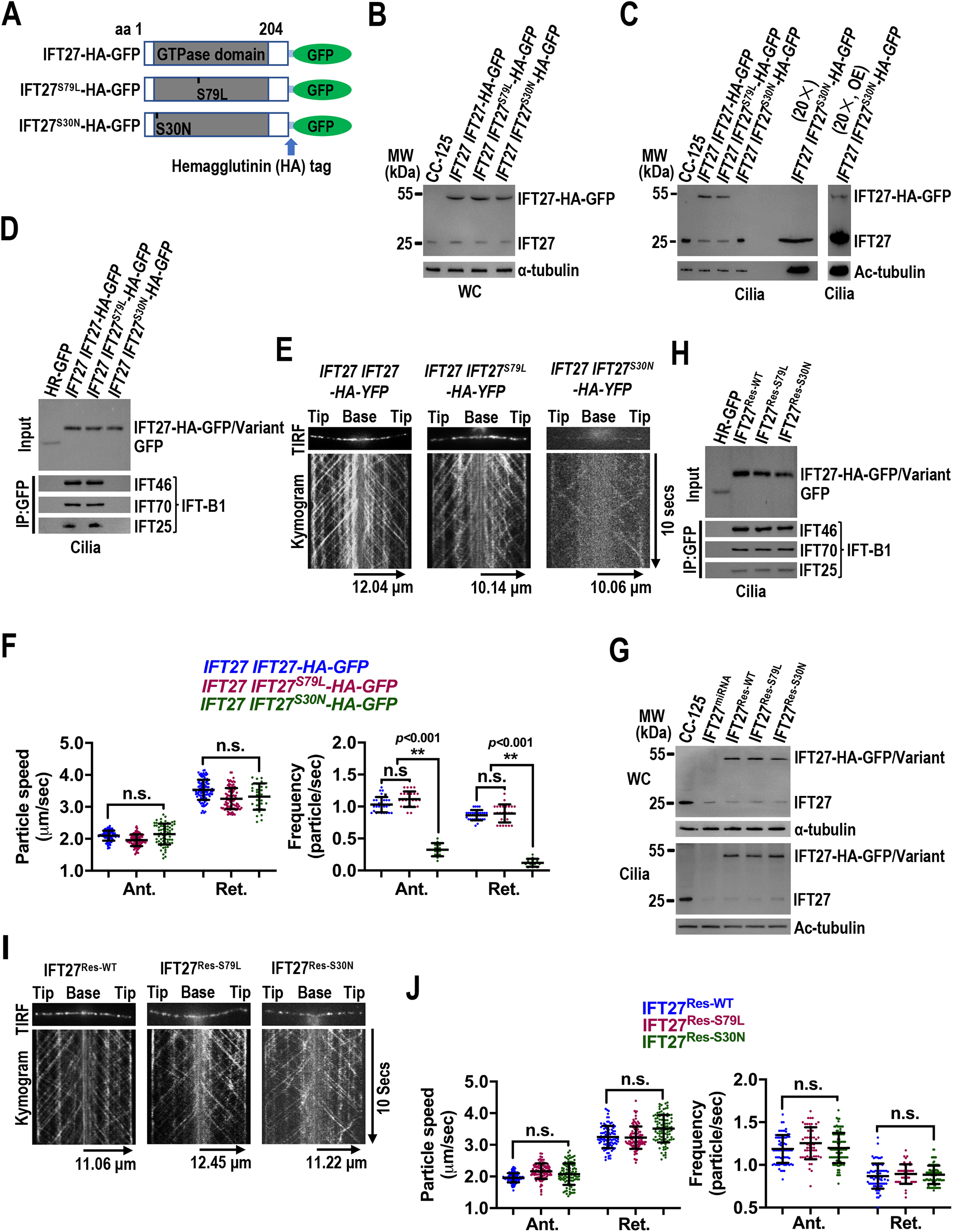
GTP loading enhances IFT27’s binding affinity for IFT-B1. (**A**) Schematic presentation of IFT27-HA-GFP, IFT27^S79L^-HA-GFP, and IFT27^S30N^-HA-GFP. The hemagglutinin (HA) tag in between the IFT27/variants and its N-terminal GFP tag was shown. (**B** and **C**) Immunoblots of WC samples (**B**) and cilia (**C**) of CC-125, *IFT27 IFT27-HA-GFP, IFT27 IFT27*^*S79L*^*-HA-GFP* and *IFT27 IFT27*^*S30N*^*-HA-GFP* cells probed with α-IFT27. Twenty times more proteins from cilia of *IFT27 IFT27*^*S30N*^*-HA-GFP* cells was also loaded and the blot was over-exposed (OE). (**D**) Immunoblots of α-GFP captured proteins from cilia of the listed cells probed for IFT46, IFT70 and IFT25. The input was adjusted with α-GFP. **(E)** TIRF images and corresponding kymograms of the listed cells (***Figure 3-videos 1-3***, 15 fps). (**F**) Speeds and frequencies of fluorescent particles to traffic inside cilia of the listed cells. (**G**) Immunoblots of WC samples and cilia of CC-125, IFT27^Res-WT^, IFT27^Res-S79L^ and IFT27^Res-S30N^ cells probed with α-IFT27. (**H**) Immunoblots of α-GFP captured proteins from cilia of the listed cells probed for IFT46, IFT70 and IFT25. The input was adjusted with α-GFP. (**I**) TIRF images and corresponding kymograms of the listed cells (***Figure 3-videos 4-6***, 15 fps). (**J**) Speeds and frequencies of the fluorescent particles to traffic inside cilia of the listed cells. For panels *B, C* and *G*, α-tubulin and Ac-tubulin were used to adjust the loading of WC samples and cilia, respectively. For panels *E* and *I*, the time and transport length are indicated on the right and on the bottom, respectively. For panels *F* and *J*, error bar indicates S.D. n.s.: non-significance. **: significance at *p*<0.001.

To examine whether the GDP-bound IFT27 can bind IFT-B1 and enter and cycle through cilia efficiently in the absence of the endogenous IFT27, we expressed IFT27^S79L^-HA-GFP and IFT27^S30N^-HA-GFP in IFT27^miRNA^ cells (resulting strains IFT27^Res-S79L^ and IFT27^Res-S30N^ cells) (***Figure 3G***). When expressed at similar amount in three transgenic cells, IFT27^S30N^-HA-GFP, similar to IFT27-HA-GFP and IFT27^S79L^-HA-GFP, entered cilia (***Figure 3G***) by binding IFT-B1 (***Figure 3H***) and underwent IFT at similar speeds and frequencies (***Figure 3I, J; Table 1; Figure 3-videos 4-6***) (Sun *et al*., 2021), suggesting that GDP-bound IFT27 can bind IFT-B1 to enter and cycle through cilia efficiently in the absence of the endogenous IFT27. Taken together, we conclude that IFT27 can bind IFT-B1 independent of its nucleotide state for ciliary entry and cycling. However, GTP loading enables IFT27 to outcompete its GDP-bound version for binding IFT-B1.

### IFT25/27 partially dissociates from IFT-B1 in cilia in a nucleotide-independent manner

Our previous studies and others have shown that IFT25 binds IFT27 to stabilize the latter (Dong *et al*., 2017b; Eguether *et al*., 2014; Liew *et al*., 2014; Wang *et al*., 2009). When expressed in bacteria alone, IFT25 is completely soluble while IFT27 is almost completely insoluble (Dong *et al*., 2017b). In contrast, the N-terminal GST-tagged IFT25 (GST-IFT25) and the N-terminal 6×His-tagged IFT27 (6×His-IFT27) both are present in the soluble fraction as determined by tandem purification performed on a GST-IFT25 and 6×His-IFT27 co-expression bacterial system (Dong *et al*., 2017b). To examine whether nucleotide state affects IFT27 binding to IFT25, we developed two bacterial systems co-expressing GST-IFT25 and 6×His-IFT27^S79L^ or GST-IFT25 and 6×His-IFT27^S30N^. Tandem purification showed that IFT27^S79L^ and IFT27^S30N^ mutants can pull down GST-IFT25 at similar amount as IFT27 does (***Figure 4A***) (Dong *et al*., 2017b). In addition, in the presence of GTPγS, GDP, or none of them, which locks IFT27 in a GTP-bound, GDP-bound, or WT state, IFT25-HA-GFP immunoprecipitated similar amounts of IFT27 and the IFT-A subunits IFT43 and IFT139, the IFT-B1 subunits IFT46 and IFT70, and the IFT-B2 subunits IFT38 and IFT57 as well in cilia of IFT25-HA-GFP-expressing *IFT25 IFT25-HA-GFP* cells (***Figure 4B***) (Dong *et al*., 2017b). Reciprocally, IFT27-HA-GFP, IFT27^S79L^-HA-GFP, and IFT27^S30N^-HA-GFP recovered IFT25 and these IFT proteins at similar amounts (***Figure 4C***). These *in vitro* and *in vivo* biochemical evidence together suggest that nucleotide state does not affect IFT27 binding to IFT25 (Liew *et al*., 2014).

**Figure 4.**
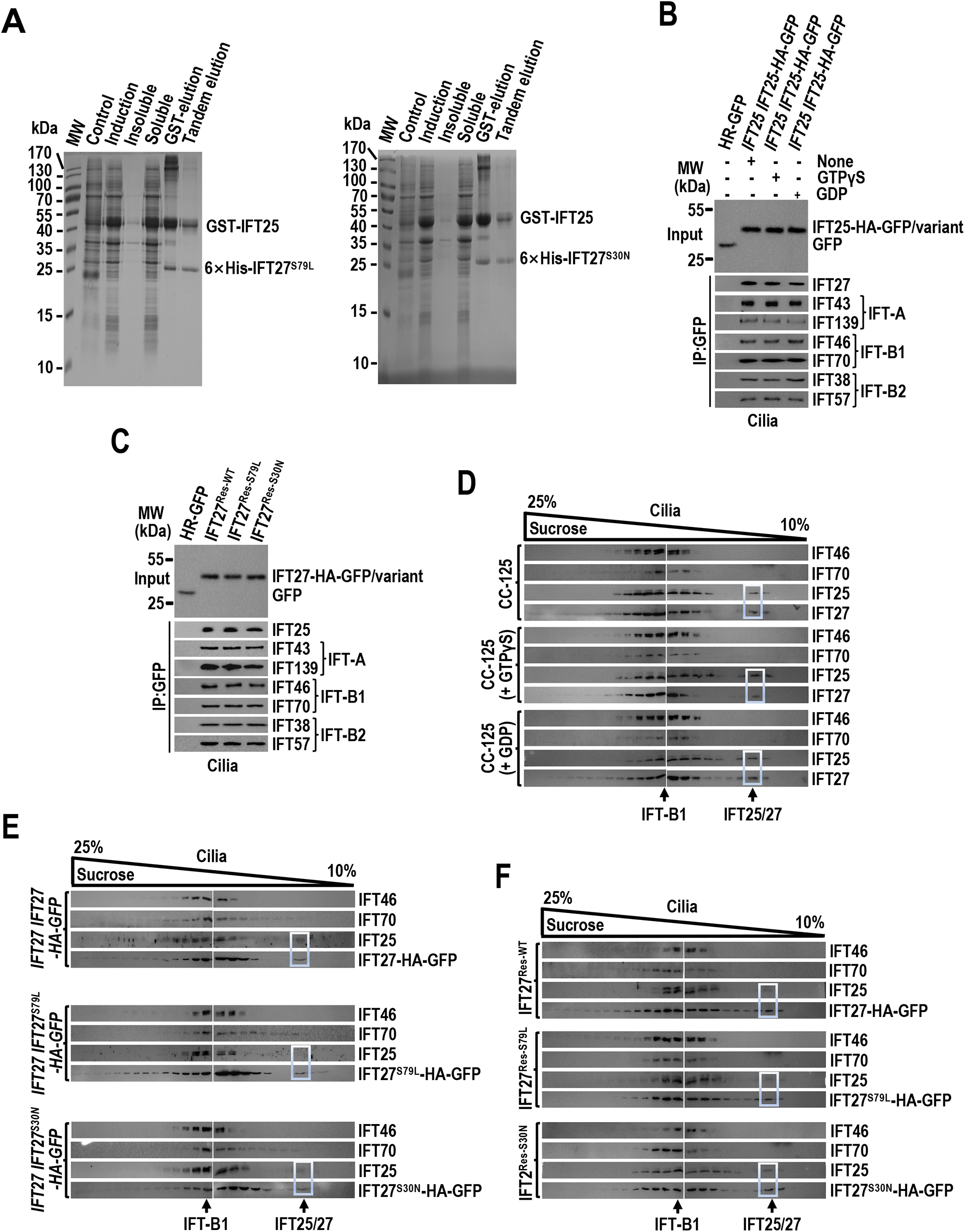
IFT25/27 partially dissociates from IFT-B1 in cilia in a nucleotide-independent manner. (**A**) Co-expression and tandem purification of recombinant GST-IFT25 and 6×His-IFT27^S79L^ (left panel) or 6×His-IFT27^S30N^ (right panel). Coomassie stained gel of samples from each step of the tandem purification; The lanes indicated as “Control” and “Induction” contained the bacterial lysates before and after IPTG induction, respectively. The lanes labeled with “Insoluble” and “Soluble” are the insoluble and soluble fractions recovered after centrifugation of cell lysates. The soluble fractions further underwent tandem purification with glutathione-agarose beads and Ni-NTA resin. (**B** and **C**) Immunoblots of α-GFP captured proteins from cilia of *IFT25 IFT25-HA-GFP* cells (**B**) and *IFT27 IFT27-HA-GFP, IFT27 IFT27*^*S79*^*-HA-GFP*, and *IFT27 IFT27*^*S30N*^*-HA-GFP* cells (**C**) probed for IFT27 (**B**) and IFT25 (**C**) and the IFT-A subunits IFT43 and IFT139, the IFT-B1 subunits IFT46 and IFT70, and the IFT-B2 subunits IFT38 and IFT57 (**B** and **C**). The input was adjusted with α-GFP by immunoblotting. (**D**) Immunoblots of sucrose density gradient of cilia of CC-125 cells in the presence of GTPγS, GDP or none of them probed for IFT46, IFT70, IFT25, and IFT27. (**E**) Immunoblots of sucrose density gradient of cilia of *IFT27 IFT27-HA-GFP, IFT27 IFT27*^*S79L*^*-HA-GFP*, and *IFT27 IFT27*^*S30N*^*-HA-GFP* cells probed for IFT46, IFT70, IFT25, and IFT27-HA-GFP or its HA-GFP-tagged variants. (**F**) Immunoblots of sucrose density gradient of cilia of IFT27^Res-WT^, IFT27^Res-S79L^, and IFT27^Res-S30N^ cells probed for IFT46, IFT70, IFT25 and IFT27-HA-GFP or its HA-GFP-tagged variants.

In agreement with our previous study, sucrose gradient density centrifugation of cilia revealed that the majority of *Chlamydomonas* IFT25/27 integrates into IFT-B1 with a minor portion also existing as a free form independent of IFT-B1 (***Figure 4D***) (Wang *et al*., 2009). The ratio of the IFT-B1-associated IFT25/27 to the IFT-B1-separated IFT25/27 likely remained unchanged in cilia in the presence of GTPγS and GDP, suggesting that the nucleotide state of IFT27 has no effect on the dissociation of IFT25/27 from IFT-B1 (***Figure 4D***). This notion was confirmed by the observation that the majority of IFT27-HA-GFP and the HA-GFP-tagged IFT27^S79L^ and IFT27^S30N^ variants co-sedimented with IFT25 and the IFT-B1 subunits IFT46 and IFT70 only with a minor same proportional portion binding IFT25 alone in cilia of IFT27-HA-GFP, IFT27^S79L^-HA-GFP and IFT27^S30N^-HA-GFP cells (***Figure 4E***). In the absence of the endogenous IFT27, similar co-sedimentation patterns of IFT25/27 were observed in cilia of IFT27^Res-WT^, IFT27^Res-S79L^ and IFT27^Res-S30N^ cells (***Figure 4F***). Thus, dissociation of IFT25/27 from IFT-B1 in cilia is unlikely controlled by IFT27 nucleotide state. Of note, TIRF imaging observed that IFT27-HA-GFP and its HA-GFP-tagged variants rarely uncouple with IFT of both anterograde and retrograde directions in cilia of IFT27^Res-WT^, IFT27^Res-S79L^ and IFT27^Res-S30N^ cells (***Figure 3I*; *Figure 3-videos 4-6***). Therefore, we propose that dissociation of IFT25/27 from IFT-B1 most likely occurs at the ciliary tip, consistent with the previous rodent cell study (Liew *et al*., 2014).

### IFT25/27 promotes BBSome reassembly at the ciliary tip independent of IFT27 nucleotide state

Our previous study has shown that, during the turnaround step at the ciliary tip, the BBSome unloads from anterograde IFT trains and disassembles followed by reassembling to form intact BBSome for reloading onto retrograde IFT trains to exit cilia (Sun *et al*., 2021). In IFT27-depleting IFT27^miRNA^ cells, the BBSome subunits remain at WT levels but build up at the ciliary tip as they are defective in reassembling to form intact BBSome entities (Dong *et al*., 2017b; Sun *et al*., 2021). To have a full view on whether nucleotide state of IFT27 affects BBSome reassembly at the ciliary tip, we determined cellular and ciliary abundance of the BBSome subunits BBS1 and BBS5 in strains IFT27^miRNA^, IFT27^Res-WT^, IFT27^Res-S79L^, and IFT27^Res-S30N^. All four strains contained BBS1 and BBS5 at WT levels (***Figure 5A***). BBS1 and BBS5 built up in IFT27^miRNA^ cilia and their ciliary retention was restored back to normal in strains IFT27^Res-WT^, IFT27^Res-S79L^, and IFT27^Res-S30N^ (***Figure 5A***). Immunostaining assays identified BBS1 and BBS5 accumulate at the tip of IFT27^miRNA^ cilia, while they were restored to WT levels at the tip of IFT27^Res-WT^, IFT27^Res-S79L^ and IFT27^Res-S30N^ cilia (***Figure 5B***). Sucrose density gradient centrifugation visualized BBS1 and BBS5 to exist as individual free forms in IFT27^miRNA^ cilia, while they become co-sedimented at BBSome fractions in IFT27^Res-WT^, IFT27^Res-S79L^, and IFT27^Res-S30N^ cilia (***Figure 5C***). These data demonstrate that IFT27 is indispensable for BBSome reassembly at the ciliary tip but in a nucleotide-independent manner.

**Figure 5.**
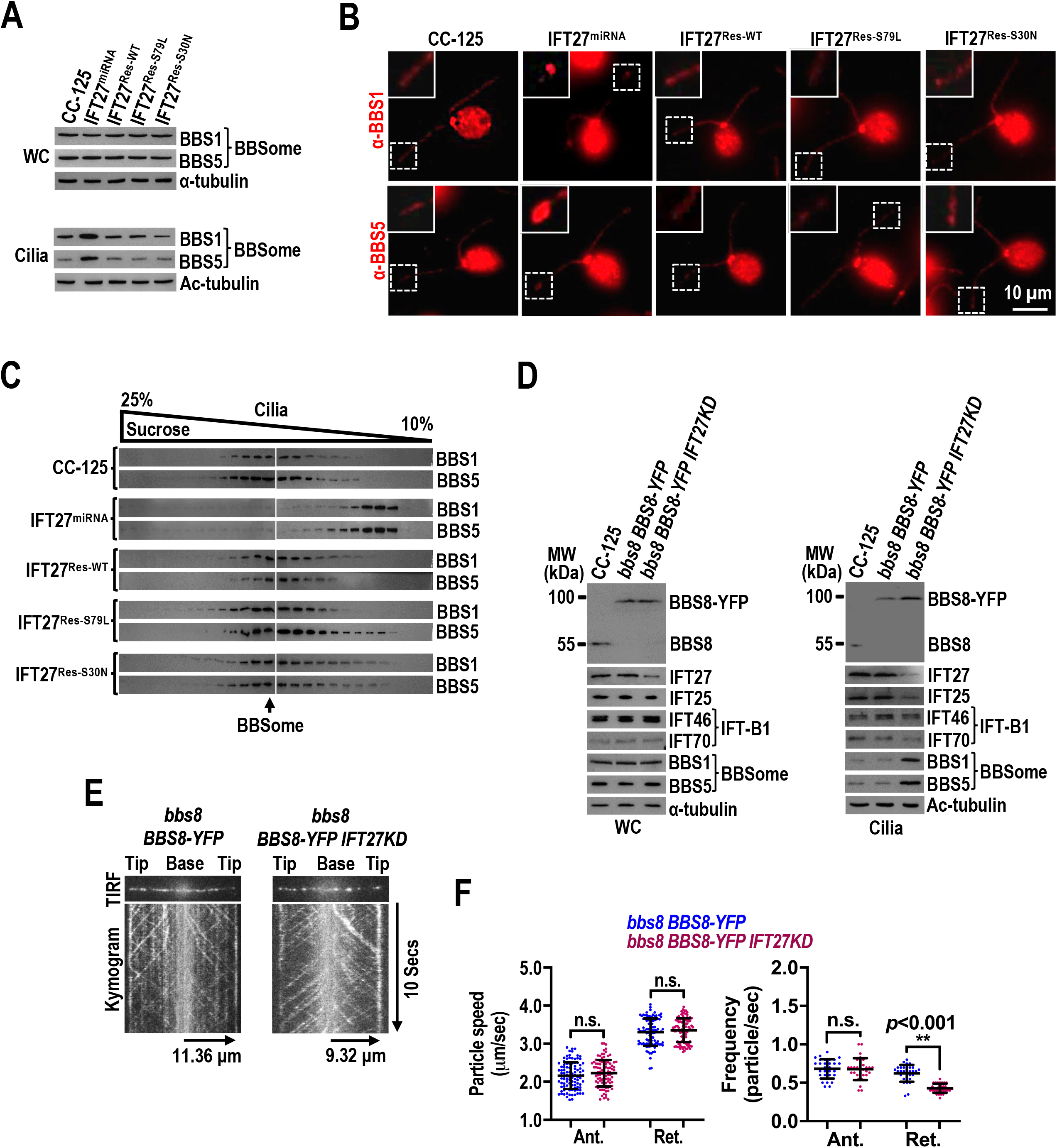
IFT27 promotes BBSome reassembly at the ciliary tip independent of its nucleotide state. (**A**) Immunoblots of WC samples and cilia of CC-125, IFT27^miRNA^, IFT27^Res-WT^, IFT27^Res-S79L^, and IFT27^Res-S30N^ cells probed for the BBSome subunits BBS1 and BBS5. Alpha-tubulin and Ac-tubulin were used to adjust the loading of WC samples and cilia, respectively. (**B**) CC-125, IFT27^miRNA^, IFT27^Res-WT^, IFT27^Res-S79L^ and IFT27^Res-S30N^ cells stained with α-BBS1 (red) and α-BBS5 (red). Inset shows the ciliary tip. Scale bars: 10 µm. (**C**) Immunoblots of sucrose density gradient of cilia of CC-125, IFT27^miRNA^, IFT27^Res-WT^, IFT27^Res-S79L^, and IFT27^Res-S30N^ cells probed with α-BBS1 and α-BBS5. (**D**) Immunoblots of WC samples and cilia of CC-125, *bbs8 BBS8-YFP*, and *bbs8 BBS8-YFP IFT27KD* cells probed with α-BBS8, α-IFT25, α-IFT27, α-IFT46, α-IFT70, α-BBS1, and α-BBS5. (**E**) TIRF images and corresponding kymograms of the listed cells (***Figure 5-videos 1*** and ***2***, 15 fps). The time and transport length are indicated on the right and on the bottom, respectively. (**F**) Speeds and frequencies of the fluorescent particles to traffic inside cilia of the listed cells. error bar indicates S.D. n.s.: non-significance. **: significance at *p*<0.001.

To visualize how the BBSome subunits accumulate at the ciliary tip in a dynamic state in the absence of IFT27, we knocked down IFT27 to ∼8% of wild-type level in the BBS8-YFP-expressing BBS8-null *bbs8 BBS8-YFP* cells, resulting strain *bbs8 BBS8-YFP IFT27KD* (***Figure 5D***) (Liu *et al*, 2021a). For *bbs8 BBS8-YFP* and *bbs8 BBS8-YFP IFT27KD* cells, BBS8-YFP, BBS1, and BBS5 were expressed at WT levels (***Figure 5D***). Once entering cilia, three BBSome components retained at WT levels in *bbs8 BBS8-YFP* cilia (***Figure 5D***). In contrast, *bbs8 BBS8-YFP IFT27KD* cilia accumulated them ∼4 folds higher than *bbs8 BBS8-YFP* cilia (***Figure 5D***). As compared to *bbs8 BBS8-YFP* cells in which BBS8-YFP undergoes bidirectional IFT at typical speed and frequency as *Chlamydomonas* BBSome subunits (Lechtreck *et al*., 2009a), *BBS8-YFP IFT27KD* cells driven BBS8-YFP to shuttle between the ciliary base and tip at normal IFT speeds and anterograde frequency but at significantly reduced retrograde frequency (***Figure 5E, F; Table 1; Figure 5-videos 1 and 2***). Besides, knockdown of IFT27 did not alter cellular content of IFT25, IFT46, and IFT70 but deprived IFT25/27 rather than IFT46 and IFT70 from cilia, leaving an intact IFT system in the absence of IFT27 (***Figure 5D***). These results together suggest that, in the absence of IFT27, the BBSome undergoes normal anterograde IFT to reach the ciliary tip but fails to exit cilia via retrograde IFT, thus accumulating at the ciliary tip.

### IFT27 mediates phototaxis via maintaining BBSome-dependent PLD ciliary removal

It has been shown that the maintenance of PLD ciliary turnover is a prerequisite for *C. reinhardtii* to perform phototaxis in response to a light stimulus (Liu & Lechtreck, 2018). PLD relies on interaction with the BBSome for loading onto retrograde IFT trains for ciliary removal (Liu & Lechtreck, 2018). In consequence, BBSome malfunction causes PLD retention in cilia, leading to non-phototactic phenotype in *C. reinhardtii* (Lechtreck *et al*., 2009a; Liu & Lechtreck, 2018). According to our current and previous studies, knockdown of IFT25 or IFT27 does not affect cellular abundance of the BBSome but causes it unable to move out of cilia so as to accumulate at the ciliary tip (Fig. 5) (Dong *et al*., 2017b; Sun *et al*., 2021). As expected, knockdown of IFT27 did not alter cellular PLD abundance but resulted in PLD accumulation in cilia as a direct result of the disrupted BBSome removal from cilia (***Figure 6A***). Following the restoration of BBSome reassembly in strains IFT27^Res-WT^, IFT27^Res-S79L^, and IFT27^Res-S30N^, PLD ciliary accumulation was rescued back to normal (***Figure 6A***). Immunostaining failed to visualize PLD ciliary distribution owing to its low ciliary abundance (Liu & Lechtreck, 2018). However, knockdown of IFT27 caused PLD to accumulate along the whole length of cilia but mostly at the ciliary tip, which can be easily visualized by immunostaining (***Figure 6B***). This ciliary PLD staining pattern was not detectable in strains IFT27^Res-WT^, IFT27^Res-S79L^, and IFT27^Res-S30N^ (***Figure 6B***). As shown by both population and single cell assays determining cell’s locomotive reaction in response to light, the IFT27^miRNA^ strain was non-phototactic (***Figure 6C, D***) (Sun *et al*., 2021). The IFT27-rescuing strain IFT27^Res-S79L^ and IFT27^Res-S30N^, like CC-125 cells, became normal in phototaxis (***Figure 6C, D***). Therefore, IFT27 mediates phototaxis via maintaining BBSome-dependent PLD removal out of cilia in *C. reinhardtii*. This process does not depend on IFT27 nucleotide state.

**Figure 6.**
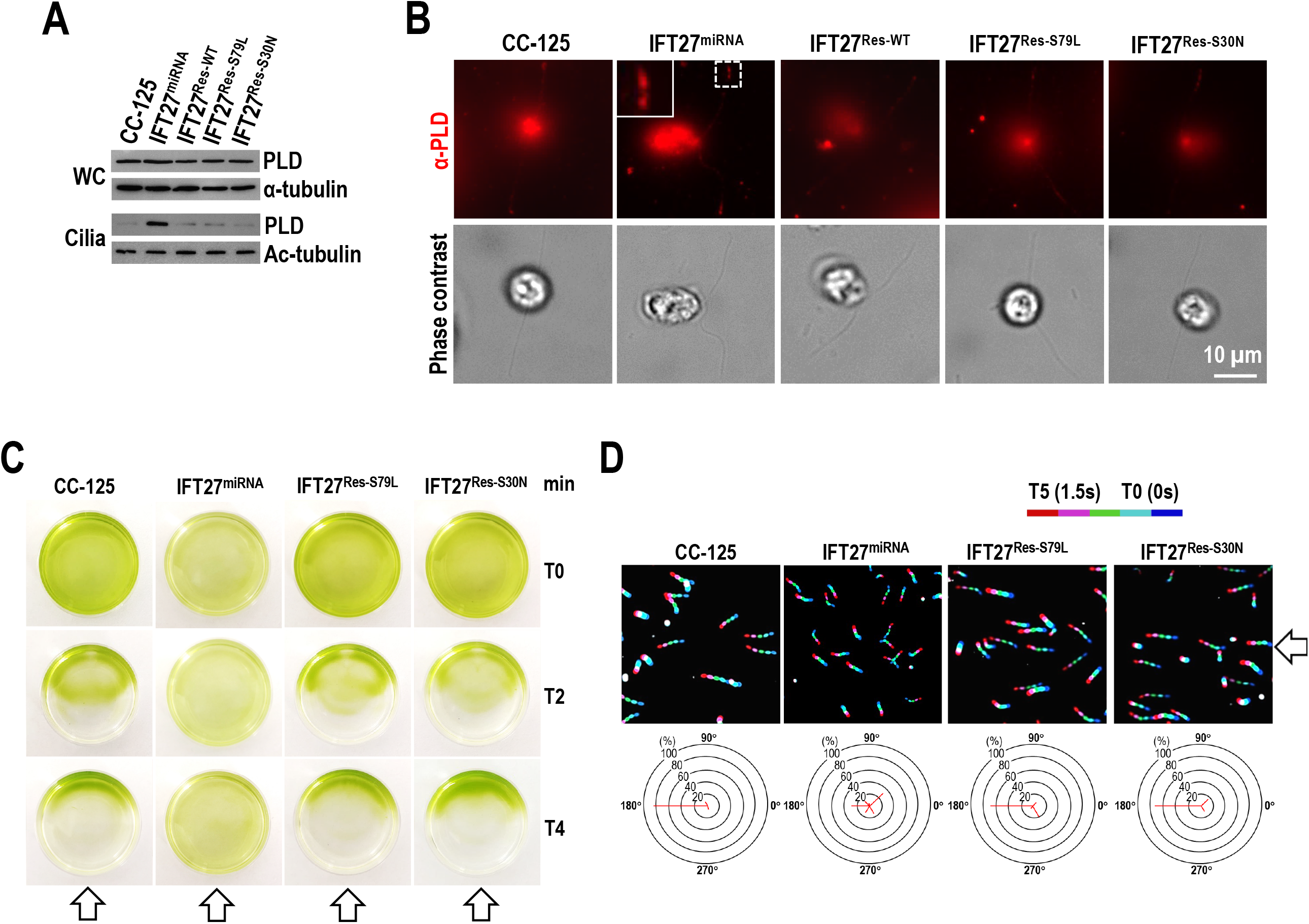
IFT27 mediates phototaxis via promoting BBSome-dependent PLD ciliary removal. (**A**) (**A**) Immunoblots of WC samples and cilia of CC-125, IFT27^miRNA^, IFT27^Res-WT^, IFT27^Res-S79L^, and IFT27^Res-S30N^ cells probed for PLD. Alpha-tubulin and Ac-tubulin were used to adjust the loading of WC samples and cilia, respectively. (**B**) CC-125, IFT27^miRNA^, IFT27^Res-WT^, IFT27^Res-S79L^ and IFT27^Res-S30N^ cells stained with α-PLD (red). Phase contrast images of the cells were shown. Inset shows the ciliary tip. Scale bars: 10 µm. (**C**) Population phototaxis assay of CC-125, IFT27^miRNA^, IFT27^Res-S79L^, and IFT27^Res-S30N^ cells. The direction of light is indicated (white arrows). (**D**) Single cell motion analysis of CC-125, IFT27^miRNA^, IFT27^Res-S79L^, and IFT27^Res-S30N^ cells. The direction of light is indicated (white arrows). The radial histograms show the percentage of cells moving in a particular direction relative to the light (six bins of 60° each). Composite micrographs showing the tracks of single cells. Each of the five merged frames was assigned a different color (blue, frame 1, and red, frame 5, corresponding to a travel time of 1.5 s). (Scale bar: 50 μm.).

## Discussion

By using *C. reinhardtii* as a model organism, we elucidated the interplay among IFT-A, IFT-B and IFT25/27, the effect of IFT27 nucleotide state on IFT27 binding to IFT25 and IFT25/27 integration into IFT-B, and the role of IFT25/27 in maintaining PLD dynamics via the BBSome for mediating phototaxis. Our data show that IFT-A, IFT-B, and IFT25/27 are irrelevant with regards to their stability. Nucleotide state of IFT27 does not affect the affinity of IFT27 for IFT25 nor IFT25/27 integration into IFT-B1. GTP-loading to IFT27 enhances the affinity of IFT25/27 for IFT-B1, while IFT25/27 cycles on and off IFT-B1 without relying on IFT27 nucleotide state. Our data also show that IFT27 promotes PLD ciliary removal via promoting BBSome reassembly at the ciliary tip and this process does not rely on nucleotide state of IFT27 (Fig. 7), closing a gap in our understanding of how IFT27 mediates phototaxis in *C. reinhardtii*.

**Figure 7.**
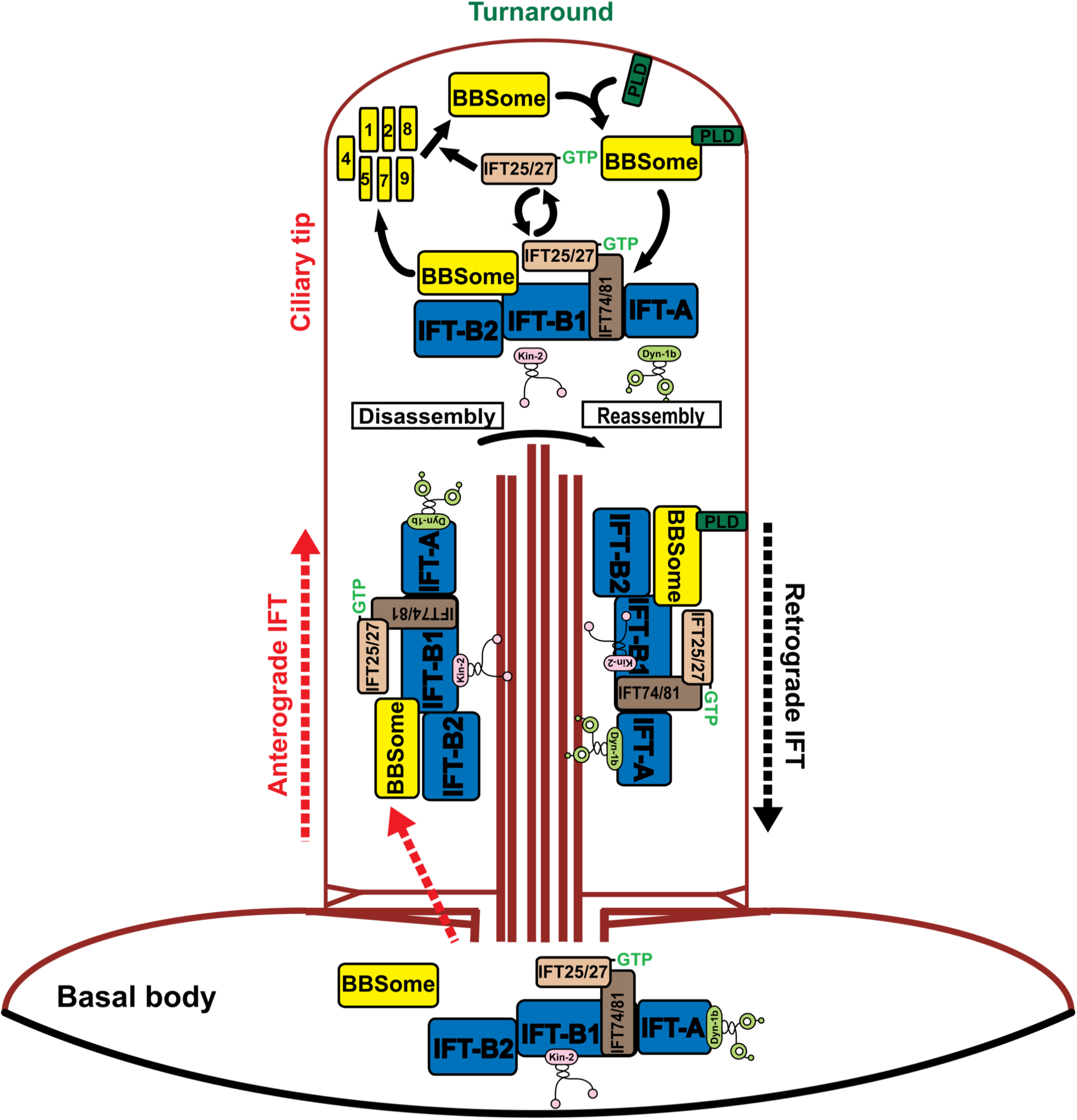
Hypothetical model how IFT25/27 mediate PLD ciliary removal via the BBSome in *C. reinhardtii*. IFT25/27-GTP binds the IFT74/81 composing component of the IFT-B1 subcomplex of IFT trains as an IFT cargo. After incorporated with the BBSome through a direct interaction between the BBSome and IFT-B1 at the basal bodies, IFT trains enter cilia for transporting to the ciliary tip by kinesin-II-mediated IFT in an anterograde direction. Upon reaching the ciliary tip, the BBSome sheds from IFT trains for remodeling. At the ciliary tip, IFT25/27-GTP can jump on and off IFT-B1 in a nucleotide-independent manner. It is the IFT train-shed IFT25/27-GTP that promotes the BBSome reassembly. The post-remodeled BBSome then interacts with the ciliary membrane anchored PLD prior to loading onto IFT trains for transporting to the ciliary base by cytoplasmic dynein 1b-mediated IFT in a retrograde direction.

### IFT25/27 is unlikely a structural component of IFT-B

As a RAB-like 4 GTPase, IFT27 binds its partner IFT25 to form the heterodimeric IFT25/27 entity cross species that harbor these two genes (Lechtreck *et al*, 2009b; Wang *et al*., 2009). IFT25 binds IFT27 to stabilize the later but not *vice versa*, suggesting that their interaction is not reciprocal (Dong *et al*., 2017b; Eguether *et al*., 2014; Keady *et al*., 2012; Sun *et al*., 2021). IFT27 nucleotide state does not affect the IFT27 affinity for IFT25, disapproving IFT27 GTPase activity in deciding IFT25 binding. Of note, GDP-loaded rather than GTP-loaded IFT27 can significantly reduce the affinity of IFT25/27 for IFT-B1 in the presence of the endogenous IFT27, suggesting that IFT27 most likely exits in a GTP-bound state in the cell (Fig. 7). This notion is supported by the facts that the cell maintains a GTP concentration at millimolar level (Woodland & Pestell, 1972), IFT25/27 binds guanine nucleotides with a micromolar affinity (Bhogaraju *et al*., 2011), and IFT27 has extremely low GTP hydrolysis activity (Bhogaraju *et al*., 2011). However, nucleotide state of IFT27 really does not influence IFT25/27 integration into IFT-B1. Of most importance, IFT-A, IFT-B, and IFT25/27 are not interdependent for maintaining their stability and loss of IFT25/27 does not impair IFT for ciliation, revealing that IFT25/27 is unlikely a structural component of IFT-B1 but rather likely binding to IFT-B1 as an IFT cargo. Other example includes RABL5/IFT22 that integrates into IFT-B1 for ciliary entry and cycling but not affects IFT for ciliation, suggesting that two Rab-like GTPase components of IFT-B1 do not play a structural role in IFT-B but a regulatory factor in cilia (Xue *et al*., 2020).

### IFT25/27 does not rely on IFT27 nucleotide state for cycling on and off IFT-B1 in cilia

Studies from both rodent cells and *C. reinhardtii* have identified IFT25/27 can uncouple with IFT-B1 at the ciliary tip (Dong *et al*., 2017b; Liew *et al*., 2014; Sun *et al*., 2021). In rodent cells, GTP hydrolysis of IFT27 confers IFT25/27 a greatly reduced affinity for IFT-B1, leading to the dynamic release of IFT25/27 from IFT-B1 at the ciliary tip (Liew *et al*., 2014). Accordingly, IFT-B1 was predicted to be an IFT27 effector and GTPase activity of IFT27 could enable IFT25/27 to cycle on and off IFT-B1 in cilia (Liew *et al*., 2014). Since this scenario was proposed based on the biochemical assays performed on whole cell samples rather than cilia itself, the proposed model may not necessarily reflect the authentic role of IFT27 nucleotide state in cilia (Liew *et al*., 2014). In *C. reinhardtii*, GTP-loaded IFT27, like its rodent counterpart (Liew *et al*., 2014), can enhance the affinity of IFT25/27 for IFT-B1 and nucleotide state of IFT27 does not affect IFT25/27 integration into IFT-B1 for ciliary entry and cycling. Upon at the ciliary tip, IFT25/27 does not rely on IFT27 nucleotide state for cycling on and off IFT-B1, results disapproving IFT-B1 to be an IFT27 effector and excluding IFT27 GTPase activity from controlling the interplay between IFT25/27 and IFT-B1. As compared to the RABL4 GTPases of other species, *Chlamydomonas* IFT27 contains a serine rather than the conserved glutamine residue in the site responsible for GTP hydrolysis and this causes IFT27 to has very low GTPase activity (Bhogaraju *et al*., 2011). We then propose that IFT25/27 composed of IFT25 and GTP-bound IFT27 (IFT25/27-GTP) binds IFT-B1 for entering and cycling through cilia. At the ciliary tip, IFT25/27-GTP likely cycles on and off IFT-B1 by an unknown mechanism (Fig. 7). Although it remains elusive whether IFT27 can hydrolyze GTP in cilia, IFT25/27 cycling on and off IFT-B1 is unlikely controlled by IFT27 GTPase activity.

### IFT25/27 does not rely on IFT27 nucleotide state for promoting BBSome ciliary tip reassembly

Mammalian IFT25 and IFT27 knock-outs accumulate the BBSome and putative BBSome cargoes at the ciliary tip, indicating that these small IFT proteins mediate BBSome loading onto retrograde IFT trains (Eguether *et al*., 2014; Keady *et al*., 2012; Liew *et al*., 2014). Our previous studies have shown that IFT25/27 is indispensable for BBSome reassembly at the ciliary tip rather than decides BBSome loading onto retrograde IFT trains for ciliary removal (Dong *et al*., 2017b; Sun *et al*., 2021). Given that IFT27^S79L^ and IFT27^S30N^ both resemble the endogenous IFT27 for promoting BBSome reassembly at the ciliary tip, nucleotide state of IFT27 is proposed not to affect the role of IFT25/27 in this process. Unlike IFT22 which enters cilia by binding IFT-B1 as its GTP-specific effector, GTP loading to IFT27 enhances the affinity of IFT25/27 to bind IFT-B1 at the basal body for ciliary entry. Upon entering cilia, IFT25/27 can release from IFT-B1 in a nucleotide-independent manner and this event most likely occurs at the ciliary tip. Therefore, it is desirable to examine whether releasing of IFT25/27 from IFT at the ciliary tip would represent any physiological roles in cilia. In addition, it has been proposed that the leucine zipper transcription factor-like 1 (LZTFL1) protein bridges the BBSome to retrograde IFT trains through IFT25/27 in cilia (Eguether *et al*., 2014). The depletion of LZTFL1 thus causes the BBSome to accumulate in cilia (Eguether *et al*., 2014; Seo *et al*, 2011). However, LZTFL1 was observed not to be able to enter cilia of some cell types (Seo *et al*., 2011). At least for *C. reinhardtii*, we have identified LZTFL1 stabilizes IFT25/27 in cytoplasm so as to control its presence and amount at the basal bodies for entering cilia (Sun *et al*., 2021). This causes BBSome retention in cilia by impairing BBSome reassembly at the ciliary tip (Sun *et al*., 2021).

### How does IFT25/27 maintain PLD dynamics in cilia for phototaxis

PLD is a negative regulator of phototaxis in *C. reinhardtii* (Liu & Lechtreck, 2018). Our previous study and others have shown that *bbs* mutants *bbs1-1, bbs4-1, bbs7-1* and *bbs8* all fail to assemble the intact BBSome in cytoplasm for ciliary entry, accumulating PLD in cilia for causing defects in phototaxis (Lechtreck *et al*., 2009a; Liu & Lechtreck, 2018; Liu *et al*., 2021a). Underlying this non-phototactic phonotype is the observation that PLD interacts with the IFT adaptor BBSome at the ciliary tip for loading onto retrograde IFT trains for ciliary export (Liu & Lechtreck, 2018; Liu *et al*., 2021b). According to this fact, any factors that disrupts BBSome assembly and ciliary entry, turnround, and export, are supposed to cause PLD accumulation in cilia, depriving *C. reinhardtii* of phototaxis. Our previous study has shown that IFT25/27, once shedding from IFT trains at the ciliary tip, can promote the BBSome reassembly (Sun *et al*., 2021). Though the mechanism behind how IFT25/27 promotes the BBSome reassembly remains unknown, this reassembly process is accomplished in an IFT27 nucleotide state-independent manner. In such a strategy, the BBSome are made available for PLD to interact with at the ciliary tip and then the PLD-laden BBSome can load onto retrograde IFT trains for ciliary retrieval, providing IFT25/27 a regulatory role in maintaining PLD dynamics in cilia (Fig. 7).

### Does IFT27 play a species- and tissue/cell-specific role in regulating ciliation and cell growth?

Other than its role in promoting BBSome reassembly during turnaround at the ciliary tip, IFT27 was reported, once depleted to ∼10.1% of WT level by RNAi interference (we performed quantification analysis based on the immunoblots as shown in the original article), to cause defects in ciliation and cell growth in *C. reinhardtii* (Qin *et al*., 2007). It was interesting to see that, different from other IFT-B1 subunits, which is generally required for stabilizing IFT-B but not IFT-A, *Chlamydomonas* IFT27, once depleted, causes disassembly of both IFT-A and IFT-B, which eventually leads to formation of short cilia and even cytokinesis defects with a yet unknown mechanism (Qin *et al*., 2007). Since a functional rescue experiment was not performed to verify the phenotype observed in the original article, the conclusion was thus suspicious. Afterwards, authority of this conclusion was further questioned as IFT25- and IFT27-null mice, IFT27-depleted mouse cells, and IFT25-knockdown *Chlamydomonas* cells all did not show IFT, ciliary, and cytokinesis defects (Dong *et al*., 2017b; Eguether *et al*., 2014; Keady *et al*., 2012; Liew *et al*., 2014). We are unable to observe any defects in ciliation and cytokinesis in *C. reinhardtii* even when IFT27 was depleted to as low as ∼8% of wild-type level by miRNA knockdown, further questioning the role of *Chlamydomonas* IFT27 in regulating ciliation and cell growth. This puzzle could be eventually clarified only when an IFT27-null mutant become available. In addition, *Trypanosoma brucei* IFT27, when depleted by RNAi interference, causes defects in ciliogenesis by disassembling IFT-B and growth defects (Huet *et al*., 2014), in line with the essential role of cilia for the trypanosome cell cycle (Kohl *et al*, 2003). Other than this example, sperm flagellum formation is found to rely on IFT25/27 in mouse (Liu *et al*, 2017; Zhang *et al*, 2017). These examples suggest species- and tissue/cell-specific differences in the role of IFT27 in ciliation and cell growth. Similarly, RABL5/IFT22 is dispensable for ciliation in mammalian and *Chlamydomonas* cells but essential for ciliation and cell growth in *Trypanosoma* adding the notion that numerous species-specific differences in the IFT machinery have developed (Takei *et al*, 2018; Wachter *et al*., 2019; Xue *et al*., 2020).

## Materials and methods

| **Key Resources Table** |  |  |  |  |
| --- | --- | --- | --- | --- |
| **Reagent type (species) or resource** | **Designation** | **Source or reference** | **Identifiers** | **Additional information** |
| genetic reagent (*C. reinhardtii*) | CC-125 | *Chlamydomonas* Resource Center (CRC) |  | Wild-type strain<br>*nit1*; *nit2*; *mt*<sup>+</sup> |
| genetic reagent (*C. reinhardtii*) | *bld1/ift52* | (Brazelton *et al.*, 2001) |  | *nit1*; *nit2*; *ift52*; *mt*<sup>+</sup> |
| genetic reagent (*C. reinhardtii*) | *bld2/ift46-1* | (Hou *et al.*, 2007) |  | *nit1*; *nit2*; *ift46*; *mt*<sup>+</sup> |
| genetic reagent (*C. reinhardtii*) | *ift43* | (Zhu *et al.*, 2017) |  | *nit1*; *nit2*; *ift43*; *mt*<sup>+</sup> |
| genetic reagent (*C. reinhardtii*) | *ift139* | (Zhu *et al.*, 2017) |  | *nit1*; *nit2*; *ift139*; *mt*<sup>+</sup> |
| genetic reagent (*C. reinhardtii*) | IFT25<sup>miRNA</sup> | (Dong *et al.*, 2017b) |  | *nit1*; *nit2*; *mt*<sup>+</sup>; IFT25KD |
| genetic reagent (*C. reinhardtii*) | IFT27<sup>miRNA</sup> | (Sun *et al.*, 2021) |  | *nit1*; *nit2*; *mt*<sup>+</sup>; IFT27KD |
| genetic reagent (*C. reinhardtii*) | IFT27<sup>Res-WT</sup> | (Sun *et al.*, 2021) |  | *nit1*; *nit2*; *mt*<sup>+</sup>; IFT27KD;<br>IFT27-HA-GFP-TG |
| genetic reagent (*C. reinhardtii*) | IFT27<sup>Res-S79L</sup> | This study |  | *nit1*; *nit2*; *mt*<sup>+</sup>; IFT27KD;<br>IFT27<sup>S79L</sup>-HA-GFP-TG |
| genetic reagent (*C. reinhardtii*) | IFT27<sup>Res-S30N</sup> | This study |  | *nit1*; *nit2*; *mt*<sup>+</sup>; IFT27KD;<br>IFT27<sup>S30N</sup>-HA-GFP-TG |
| genetic reagent (*C. reinhardtii*) | IFT43 IFT43-HA-YFP | This study |  | *nit1*; *nit2*; *mt*<sup>+</sup>; IFT43-HA-YFP-TG |
| genetic reagent (*C. reinhardtii*) | IFT43 IFT43-HA-YFP IFT27KD | This study |  | *nit1*; *nit2*; *mt*<sup>+</sup>; IFT27KD;<br>IFT43-HA-YFP-TG |
| genetic reagent (*C. reinhardtii*) | IFT27 IFT27-HA-GFP | This study |  | *nit1*; *nit2*; *mt*<sup>+</sup>; IFT27-HA-GFP-TG |
| genetic reagent (*C. reinhardtii*) | IFT27 IFT27<sup>S79L</sup>-HA-GFP | This study |  | *nit1*; *nit2*; *mt*<sup>+</sup>; IFT27<sup>S79L</sup>-HA-GFP-TG |
| genetic reagent (*C. reinhardtii*) | IFT27 IFT27<sup>S30N</sup>-HA-GFP | This study |  | *nit1*; *nit2*; *mt*<sup>+</sup>; IFT27<sup>S30N</sup>-HA-GFP-TG |
| genetic reagent (*C. reinhardtii*) | IFT25 IFT25-HA-GFP | (Dong *et al.*, 2017b) |  | *nit1*; *nit2*; *mt*<sup>+</sup>; IFT25-HA-GFP-TG |
| genetic reagent (*C. reinhardtii*) | *bbs8* BBS8-YFP | (Liu *et al.*, 2021a) |  | *bbs8*; *mt*<sup>+</sup>; BBS8-YFP-TG |
| genetic reagent (*C. reinhardtii*) | *bbs8* BBS8-YFP IFT27KD | This study |  | *bbs8*; *mt*<sup>+</sup>; IFT27KD;<br>BBS8-YFP-TG |
| genetic reagent (*C. reinhardtii*) | HR-GFP | (Dong *et al.*, 2017a) |  | *nit1*; *nit2*; *mt*<sup>+</sup>; HA-GFP-TG |
| antibody | Anti-IFT25 (rabbit polyclonal) | (Dong *et al.*, 2017b) |  | immunoblotting (1:500) |
| antibody | Anti-IFT27 (rabbit polyclonal) | (Dong *et al.*, 2017b) |  | immunoblotting (1:500) |
| antibody | Anti-IFT38 (rabbit polyclonal) | (Sun *et al.*, 2021) |  | immunoblotting (1:1,000) |
| antibody | Anti-IFT43 (rabbit polyclonal) | (Zhu *et al.*, 2017) |  | immunoblotting (1:500) |
| antibody | Anti-IFT46 (rabbit polyclonal) | (Dong *et al.*, 2017b) |  | immunoblotting (1:1,000) |
| antibody | Anti-IFT57<br>(rabbit polyclonal) | (Dong *et al.*, 2017b) |  | immunoblotting (1:500) |
| antibody | Anti-IFT70<br>(rabbit polyclonal) | (Dong *et al.*, 2017b) |  | immunoblotting (1:1,000) |
| antibody | Anti-IFT139<br>(rabbit polyclonal) | (Dong *et al.*, 2017b) |  | immunoblotting (1:1,000) |
| antibody | Anti-FLA10<br>(rabbit polyclonal) | (Dong *et al.*, 2017b) |  | immunoblotting (1:1,000) |
| antibody | Anti-D1bLIC<br>(rabbit polyclonal) | (Dong *et al.*, 2017b) |  | immunoblotting (1:1,000) |
| antibody | Anti-BBS1<br>(rabbit polyclonal) | (Xue *et al.*, 2020) |  | immunoblotting (1:1,000)<br>immunofluorescence (1:200) |
| antibody | Anti-BBS5<br>(rabbit polyclonal) | (Xue *et al.*, 2020) |  | immunoblotting (1:1000)<br>immunofluorescence (1:200) |
| antibody | Anti-BBS8<br>(rabbit polyclonal) | (Sun *et al.*, 2021) |  | immunoblotting (1:500) |
| antibody | Anti-PLD<br>(rabbit polyclonal) | (Liu *et al.*, 2021b) |  | immunoblotting (1:1000)<br>immunofluorescence (1:200) |
| antibody | Anti-α-tubulin<br>(mouse monoclonal) | Sigma-Aldrich | B512 | immunoblotting (1:25,000) |
| antibody | Anti-acetylated-tubulin<br>(mouse monoclonal) | Sigma-Aldrich | 6-11B-1 | immunoblotting (1:50,000) |
| antibody | Anti-GFP<br>(mouse monoclonal) | Roche | 7.1 and<br>13.1 | immunoblotting (1:1,000) |
| antibody | HRP-conjugated<br>goat anti-mouse<br>IgG | The Jackson Lab. | RRID:<br>AB_10015<br>289 | immunoblotting (1:10,000) |
| antibody | HRP-conjugated<br>goat anti-rabbit<br>IgG | The Jackson Lab. | RRID:<br>AB_23135<br>67 | immunoblotting (1:10,000) |
| antibody | Alexa-Fluor 594-conjugated goat<br>anti-rabbit IgG | Molecular Probes | RRID:<br>AB_14205<br>7 | immunofluorescence (1:400) |

### Chlamydomonas strains and growth conditions

*C. reinhardtii* strains CC-125, IFT52-null *bld1/ift52* (Brazelton *et al*., 2001), and IFT46-null *bld2/ift46-1* (Hou *et al*., 2007) mutants were bought from the *Chlamydomonas* Resource Center at the University of Minnesota, Twin Cities, MN (http://www.chlamy.org). IFT43-null *ift43* and IFT139-null *ift139* mutants were gifts from Dr. Junming Pan at Tsinghua University (Zhu *et al*., 2017). IFT25- and IFT27-knockdown strains IFT25^miRNA^ and IFT27^miRNA^, IFT27-rescuing strain IFT27^Res-WT^, *bbs8 BBS8- YFP*, and GFP-expressing strain HR-GFP have been reported previously (Dong *et al*., 2017a; Dong *et al*., 2017b; Liu *et al*., 2021a; Sun *et al*., 2021). The transgenic strains including *IFT27 IFT27-HA-GFP, IFT27 IFT27*^*S79L*^*-HA-GFP, IFT27 IFT27*^*S30N*^*-HA-GFP*, IFT27^Res-S79L^, IFT27^Res-S30N^, *IFT43 IFT43-HA-YFP, IFT43 IFT43-HA-YFP IFT27KD*, and *bbs8 BBS8-YFP IFT27KD* were generated as describe below. All strains were listed in the key resources table. Tris acetic acid phosphate (TAP) medium was used to grow strains in a continuous light with constant aeration at room temperature. In a strain-specific manner, cells were grown with or without the addition of 20 µg/ml paromomycin (Sigma-Aldrich), 15 µg/ml bleomycin (Invitrogen), or both antibiotics with 10 µg/ml paromomycin and 5 µg/ml bleomycin.

### Constructs and strain generation

To express IFT27^S79L^-HA-GFP and IFT27^S30N^-HA-GFP, the 2-kb IFT27-expressing fragment was cut from pBKS-gIFT27-HA-GFP-Paro (Sun *et al*., 2021) and inserted into the *Spe*I and *Eco*RI sites of pBluescript II KS(+) vector. The desiring mutations including S79L and S30N were introduced by site-directed mutagenesis using the primer pairs IFT27S79L-FOR (5’-GCTGGACACGGCGGGGCTCGAC-CTGTACAAGGAG-3’) and IFT27S79L-REV (5’-CTCCTTGTACAGGTCGAGCCCCGCCGTGTCCA-GC-3’) and IFT27S30N-FOR (5’-GCGACTGTCGGCAAGAACGCGCTCATCTCTATG-3’) and IFT27S30N-REV (5’-CATAGAGATGAGCGCGTTCTTGCCGACAGTCGC-3’). The mutated DNAs were inserted into the *Spe*I and *Eco*RI sites of pBKS-gIFT27-HA-GFP-Paro, resulting in pBKS-gIFT27^S30N^-HA-GFP-Paro and pBKS-gIFT27^S79L^-HA-GFP-Paro. To express the rescue vector with a *Ble* resistance, the Ble cassette was digested from pBKS-gIFT25-HA-GFP-Ble (Dong *et al*., 2017b) with *Apa*I and *Kpn*I and inserted into the *Apa*I and *Kpn*I sites of the expression vector to replace the Paro cassette, resulting in pBKS-gIFT27^S30N^-HA-GFP-Ble and pBKS-gIFT27^S79L^-HA-GFP-Ble. To express IFT43-HA-YFP, a 2.8-kb IFT43 fragment consisting of 1-kb promoter sequence and its coding region was amplified from the genomic DNA with primer pair IFT43-FOR (5’-ACGCGGCCGCATGGTGTCC-TACCTGGGC-3’) and IFT43-REV (5’-GTGAATTCCAGCTTGATGGGCATGG-3’) with *Not*I and *Eco*RI restriction enzyme sites located in its 5’- and 3’-end, respectively. The fragment of HA-YFP-Ble was cut from pBKS-gBBS3-HA-YFP-Ble by *Eco*RI and *Kpn*I (Xue *et al*., 2020). Two fragments were then inserted into the *Not*I and *Kpn*I sites of pBluescript II KS(+) vector by three-way ligation, resulting in pBKS-gIFT43-HA-YFP-Ble. After these vectors were verified by direct nucleotide sequencing, they were transformed into *C. reinhardtii* strain by electroporation for positive transformant screening according to the method described previously (Xue *et al*., 2020). To generate the *bbs8 BBS8-YFP IFT27KD* strain, the IFT27 miRNA vector pMi-IFT27 (Sun *et al*., 2021) was transformed into *bbs8 BBS8-YFP* cells (Liu *et al*., 2021a) and miRNA strain screening was initiated by checking the cellular level of IFT27 through immunoblotting of whole cell extracts with IFT27 antibody (Sun *et al*., 2021).

#### DNA preparation and messenger RNA quantification

*Chlamydomonas* genomic DNA was prepared according to our published method (Dong *et al*., 2017b). To amplify IFT43 genomic sequence, 20 ng of genomic DNA was used as template and the PCR reaction was performed at 95 °C for 2 min followed by 30 cycles of 95 °C for 10 sec, 61 °C for 30 sec, and 72 °C for 4 min with the primer pair gIFT43-FOR and gIFT43-REV. IFT protein mRNAs were quantified according to our published method (Dong *et al*., 2017b). In brief, two micrograms of RNA from each sample were reverse transcribed at 42 °C for 1 h followed by 95 °C for 10 min M-MLV Reverse Transcriptase (Promega) and oligo(T)18 primers (Takara). Relative transcript amounts were measured by SYBR-green quantitative PCR (qPCR) using primer pairs that spans either introns or exons of the target genes with an a StepOnePlus Real-Time PCR System (Applied Biosystems). The PCR reactions were performed at 95 °C for 10 min followed by 45 cycles of 95 °C for 15 sec, 60 °C for 20 sec and 72 °C for 45 sec. Samples were normalized using guanine nucleotide-binding protein subunit beta-like protein (GBLP) as a housekeeping gene internal control. For each gene, mRNA was quantified with at least three repeats. The expression level was presented using GraphPad Prism 8.0 (GraphPad Software) and presented as mean ± S.D. Primer pairs used: 5’-AAGTGCGACCAGGACAAGC-3’ and 5’-GTTGAC-GGTGGCAAACTCG-3’ for *ift25*; 5’-TCTCGGTGGAGCTCTTTCTG-3’ and 5’-GCTCACATCGAACA-CGAGAA-3’ for *ift27*; 5’-GGCGGACTGCTTGTATTG-3’ and 5’-CCTTTGAGGTCGCTTTGTA-3’ for *ift38*; 5’-CAGCAAGAAGTTTGTGGGCGTGTCG-3’ and 5’-CCATGGCGGGCAGCTTGTAGTG-3’ for *ift43*; 5’-ACCGTCAACTACAGCAAGCG-3’ and 5’-GGTCGTCGTACACGGGAATG-3’ for *ift46*; 5’-G-GCGGTCCAGTAGCTGTAGGAAGGTC-3’ and 5’-GGCACGTCAAGTTCGGGTCATCAAA-3’ for *ift57*; 5’-GATTGCGTTCAACTTATCGTCC-3’ and 5’-GGTACTTGTAGTGCGTGGTGC-3’ for *ift70*; 5’-TGCTGGCTGACATCCACATAC-3’ and 5’-GCCTGGTTCT CGTGCTTCC-3’ for *ift139*; 5’-GTCATC-CACTGCCTGTGCTTCT-3’ and 5’-GGCCTTCTTGCTGGTGATGTT-3’ for GBLP.

### Cilia preparation

Method for isolating cilia has been described previously (Wang *et al*., 2009). Briefly, cells were suspended in 150 ml of TAP (pH7.4) and incubated for 2∼4 h under strong light with bubbling. Then, 0.5 M acetic acid was added to reach a pH value of 4.5 to deciliate cells before 0.5 M KOH was added to reach a pH value of 7.4. After centrifugation (600 *×g* at 4°C for 5 min), cell bodies (pellets) and cilia (supernatants) were collected separately. Cilia were then repeatedly washed with HMDEKN buffer (30 mM Hepes [pH 7.4], 5 mM MgSO_4_, 1 mM DTT, 0.5 mM EGTA, 25 mM KCl, 125 mM NaCl) by centrifugation (12,000 ×*g* at 4°C for 10 min) until the green color disappeared completely.

### Antibodies and immunoblotting

All the rabbit-raised polyclonal antibodies have been described previously (Dong *et al*., 2017b; Liu *et al*., 2021b; Sun *et al*., 2021; Xue *et al*., 2020; Zhu *et al*., 2017) (key resources table). The monoclonal antibodies against GFP (mAbs 7.1 and 13.1, Roche), α-tubulin (mAb B512, Sigma) and acetylated-tubulin (mAb 6-11B-1, Sigma-Aldrich) were commercially bought (key resources table). Preparation of whole cell and ciliary extracts, SDS-PAGE electrophoresis, immunoblotting were performed as detailed previously (Xue *et al*., 2020). If not otherwise specified, 20 µg of total protein from each sample was loaded for SDS-PAGE electrophoresis. In immunoblotting assays, a dilution used for primary and secondary antibodies was listed in the key resources table. Quantification of the target proteins was performed by measuring the intensity of the immunoblots with ImageJ software (version 1.42g, National Institutes of Health) as described previously (Xue *et al*., 2020). The intensity of immunoblots was normalized to the intensity of a loading control protein. The data was processed with Prism 8.0 (GraphPad Software) and presented as mean ± S.D.

### Immunoprecipitation

Cilia isolated from *Chlamydomonas* strains including *IFT27 IFT27-HA-GFP, IFT27 IFT27*^*S79L*^*-HA-GFP, IFT27 IFT27*^*S30N*^*-HA-GFP*, IFT27^Res-WT^, IFT27^Res-S79L^, IFT27^Res-S30N^, *IFT25 IFT25-HA-GFP*, and HR-GFP were resuspended in HMEK (30 mM Hepes, pH 7.4, 5 mM MgSO4, 0.5 mM EGTA, and 25 mM KCl) plus protein inhibitors (1 mM PMSF, 50 µg/ml soybean trypsin inhibitor, 1 µg/ml pepstatin A, 2 µg/ml aprotinin, and 1 µg/ml leupeptin) supplemented with 50 mM NaCl and lysed by adding nonidet P-40 (NP-40) to 1%. The supernatants were collected by centrifugation (14,000 ×*g*, 4 °C, 10 min) and were incubated with agitation with 5% BSA-pretreated camel anti-GFP (YFP) antibody-conjugated agarose beads (V-nanoab Biotechnology) for 2 hrs at 4 °C. The beads were then washed with HMEK containing 150 mM NaCl, 50 mM NaCl and finally 0 mM NaCl. The beads were then added with Laemmli SDS sample buffer and boiled for 5 min before centrifuging at 2,500 *×g* for 5 min. The supernatants were then analyzed by immunoblotting as described above. If necessary, the assay was performed in the presence of GTPγS (20 mM) or GDP (20 mM), respectively.

### Immunofluorescence

The details for immunofluorescence staining has been described in our previous report (Wang *et al*., 2009). The primary antibodies against BBS1, BBS5, and PLD and the secondary antibodies Alexa-Fluor594 conjugated goat anti-rabbit (Molecular Probes, Eugene, OR) were listed in the key resource table with their suggested dilutions. Images were captured with an IX83 inverted fluorescent microscopy (Olympus) equipped with a back illuminated scientific CMOS camera (Prime 95B, Photometrics) at 100**×** amplification and processed with CellSens Dimension (version 2.1, Olympus).

### Sucrose density gradient centrifugation of ciliary extracts

The details for sucrose density gradient centrifugation has been reported previously (Wang *et al*., 2009). Linear 12 ml of 10-25% sucrose density gradients in 1×HMDEKN buffer plus protease inhibitors (1 mM PMSF, 50 μg/ml soybean trypsin inhibitor, 1 μg/ml pepstatin A, 2 μg/ml aprotinin, 1 μg/ml leupeptin) and 1% NP-40 were generated by using the Jule Gradient Former (Jule, Inc. Milford) and used within 1 hour. Cilia were opened by liquid nitrogen for three rounds of frozen-and-thaw cycles and centrifuged at 12,000 *×g*, 4 °C, for 10 min. 700 micro-litter of ciliary extracts were then loaded on the top of the gradients and separated at 38,000 *×g*, 4°C, for 14 hours in a SW41Ti rotor (Beckman Coulter). The gradients were fractioned into 24 to 26 0.5 ml aliquots by using a Pharmacia LKB Pump P-1 coupled with a FRAC-100 fraction collector. The standards used to calculate S-values were BSA (4.4S), aldolase (7.35S), catalase (11.3S), and thyroglobulin (19.4S). Twenty micro-litter of each fraction was analyzed by immunoblotting as described above. If necessary, the assay was carried out in the presence of GTPγS (20 mM) or GDP (20 mM), respectively.

### IFT video imaging and speed measurements

The motility of GFP- and YFP-tagged proteins in cilia was imaged at ∼15 frames per second (fps) using total internal reflection fluorescence (TIRF) microscopy on an inverted microscope (IX83, Olympus) equipped with a through-the-objective TIRF system, a 100×/1.49 NA TIRF oil immersion objective (Olympus), and a back illuminated scientific CMOS camera (Prime 95B, Photometrics) as detailed previously (Xue *et al*., 2020). To quantify IFT speeds and rates, kymogram were generated and measured with CellSens Dimension (version 2.1, Olympus).

### Tandem purification of bacterial expressed proteins

A dual expression vector was created on a pGEX2T backbone (GE Healthcare) and two plasmids were generated for co-expression of IFT25 and IFT27 or its mutants (IFT27^S79L^ and IFT27^S30N^) (Dong *et al*., 2017b). IFT25 and IFT27 mutants were expressed with N-terminal GST (GST-IFT25) or 6×His (6×His-IFT27^S79L^ or 6×His-IFT27^S30N^) tags. Bacterial expression of the recombinant proteins has been described previously (Fan *et al*., 2010). Clarified cell extract was applied to a 0.4 ml glutathione-agarose beads (GST-IFT25) and Ni-NTA resin (6×His-IFT27^S79L^ and 6×His-IFT27^S30N^) for tandem purification of GST-IFT25 and 6×His-IFT27^S79L^ or 6×His-IFT27^S30N^ according to our previous report (Dong *et al*., 2017b). Aliquots (8 μl) from each step of the purification were resolved on 10% SDS-PAGE gels and visualized with Coomassie Blue staining.

### Small GTPase assay

Small GTPase assay was performed according to our published method (Xue *et al*., 2020). The intrinsic GTP hydrolysis of the tandemly purified IFT25/27, IFT25/27^S79L^ and IFT25/27^S30N^ was measured by an optical assay for the release of inorganic phosphate with reagents from the QuantiChrom™ ATPase/GTPase assay kit (Bioassay Systems) (Pan *et al*, 2006).

### Population phototaxis assay

Population phototaxis assays were performed on *Chlamydomonas* cells according to the protocol reported previously (Liu & Lechtreck, 2018). In brief, 100 μl of the *Chlamydomonas* cells at concentration of ∼10^7^ were placed into the surface of Petri dishes of 3.5-cm diameter (706001; Wuxi NEST Biotech.) containing TAP medium. Afterthen, the cells were illuminated with a flashlight from one side for 4 min. Images were continuously taken once every two minutes with a standard digital camera (Nikon A70).

### Single-cell motion assay

Single-cell motion assay was performed on *Chlamydomonas* cells according to the protocol reported previously (Liu & Lechtreck, 2018). Briefly, 20 μl of cell suspensions were placed on superfrost™ plus microscope slides (12-550-15; Fisher Brand) and observed using an inverted microscope (IX83, Olympus) under non-phototactic red light illumination. The cells were then illuminated with phototactic active flashlight for 5s. After that, five images were continuously taken every 0.3s using a scientific CMOS camera (Prime 95B, Photometrics). The five sequential images, displayed in different colors, were merged in ImageJ (version 1.42g, National Institutes of Health) to show the swimming tracks of single cells, allowing us to determine the angle and the direction of a cell’s movements. Excel

### Statistical analysis

Statistical analysis was performed with GraphPad Prism 8.0 (GraphPad Software). For comparisons on velocities and frequencies of the GFP- and YFP-labeled proteins, one-sample unpaired student *t*-test was used on samples from 10 cells, if not otherwise specified. The data were presented as mean ± S.D. Significance was setup at *P*<0.001 (marked as **). n.s. represents non-significance.

## Acknowledgments

This work was supported by National Natural Science Foundation of China (32070698 to Z-C.F. and 32100541 to B.X.) and China Postdoctoral Science Foundation (2021M702457 to Y-X.L. and 2021M692403 to B.X.). The founders have no role in study design, data collection and analysis, decision to publish, or preparation of the manuscript.

## Author contributions

Yan-Xia Liu, Data curation, Formal analysis, Project administration, Methodology, Investigation, Visualization, Validation; Bin Xue, Investigation, Validation; Zhen-Chuan Fan, Conceptualization, Data curation, Resources, Supervision, Funding acquisition, Writing-original draft, Project administration, Writing-review and editing, Validation

## Data availability

All data generated or analyzed during this study are included in the manuscript and supporting files.

## Figure supplement legends

**Figure 3-figure supplement 1. IFT27 is highly conserved proteins across ciliated species**. (A) Sequence alignment of deduced amino acid sequences from ten invertebrate and vertebrate IFT27 orthologues and modified IFT27 sequences. Alignment was generated using CLC main workbench (version 6.8); the most conserved residues are shown in blue, the less conserved are in red. IFT27 contains five conserved domains including G1, G2, G3, G4 and G5 as labeled in the boxes. The P-loop serine (S30) and serine (S79) were marked. Dashes indicate gaps introduced to optimize the alignment. Arrowheads indicate missense mutations created in dominant-negative and constitutive-active IFT27 mutants. The IFT27 accession numbers are *Bos Taurus*, AAI02615.1, *Ovis aries*, NP_001119838.1, *Homo sapiens*, NP_073614.1, *Macaca mulatta*, AFH27541.1, *Mus musculus*, AAH09150.1, *Rattus norvegicus*, NP_001123967.1, *Callorhinchus milii*, NP_001279585.1, *Xenopus tropicalis*, NP_001011271.1, *Danio rerio*, NP_001017884.1, and *Chlamydomonas reinhardtii*, XP_001689669.1. (B) GTP hydrolysis of IFT25/27, IFT25/27^S79L^ and IFT25/27^S30N^. Introduction of a leucine at the position of S79 of IFT27 dramatically reduces the intrinsic GTPase activity, indicating that IFT27^S79L^ is a GTP-locked constitutive-active variant. No GTPase activity was observed for IFT27^S30N^, indicating that IFT27^S30N^ appears to be an inactive GDP-locked or simply a dominant-negative variant.

## Video legends

**Figure 2-Video 1**. TIRF imaging of IFT43-HA-YFP movement in cilia of CC-125 cells (*IFT43 IFT43-HA-YFP* cells). A frame from this movie and kymogram are shown in Figure 2E. Play speed is real-time (15 fps).

**Figure 2-Video 2**. TIRF imaging of IFT43-HA-YFP movement in cilia of IFT27^miRNA^ cells (*IFT43 IFT43-HA-YFP IFT27KD* cells). A frame from this movie and kymograph are shown in Figure 2E. Play speed is real-time (15 fps).

**Figure 3-Video 1**. TIRF imaging of IFT27-HA-GFP movement in cilia of CC-125 cells (*IFT27 IFT27-HA-GFP* cells). A frame from this movie and kymograph are shown in Figure 3E. Play speed is real-time (15 fps).

**Figure 3-Video 2**. TIRF imaging of IFT27^S79L^-HA-GFP movement in cilia of CC-125 cells (*IFT27 IFT27*^*S79L*^*-HA-GFP* cells). A frame from this movie and kymograph are shown in Figure 3E. Play speed is real-time (15 fps).

**Figure 3-Video 3**. TIRF imaging of IFT27^S30N^-HA-GFP movement in cilia of CC-125 cells (*IFT27 IFT27*^*S30N*^*-HA-GFP* cells). A frame from this movie and kymograph are shown in Figure 3E. Play speed is real-time (15 fps).

**Figure 3-Video 4**. TIRF imaging of IFT27-HA-GFP movement in cilia of IFT27^miRNA^ cells (IFT27^Res-WT^ cells). A frame from this movie and kymograph are shown in Figure 3I. Play speed is real-time (15 fps).

**Figure 3-Video 5**. TIRF imaging of IFT27^S79L^-HA-GFP movement in cilia of IFT27^miRNA^ cells (IFT27^Res-S79L^ cells). A frame from this movie and kymograph are shown in Figure 3I. Play speed is real-time (15 fps).

**Figure 3-Video 6**. TIRF imaging of IFT27^S30N^-HA-GFP movement in cilia of IFT27^miRNA^ cells (IFT27^Res-S30N^ cells). A frame from this movie and kymograph are shown in Figure 3I. Play speed is real-time (15 fps).

**Figure 5-Video 1**. TIRF imaging of BBS8-YFP movement in cilia of *bbs8* cells (*bbs8 BBS8-YFP* cells). A frame from this movie and kymograph are shown in Figure 5E. Play speed is real-time (15 fps).

**Figure 5-Video 2**. TIRF imaging of BBS8-YFP movement in cilia of *bbs8 BBS8-YFP IFT27KD* cells. A frame from this movie and kymograph are shown in Figure 5E. Play speed is real-time (15 fps).

## Source data legends

**Figure 3-source data 1**. Source data for the GTP hydrolysis experiment shown in Figure 3-figure supplement 1B.

