## Supplementary figures and images for "*Chlamydomonas* RABL4/IFT27 Mediates Phototaxis via Promoting BBSome-dependent Ciliary Export of Phospholipase D"

### Figure 3-figure supplement 1

#### Figure 3-figure supplement 1

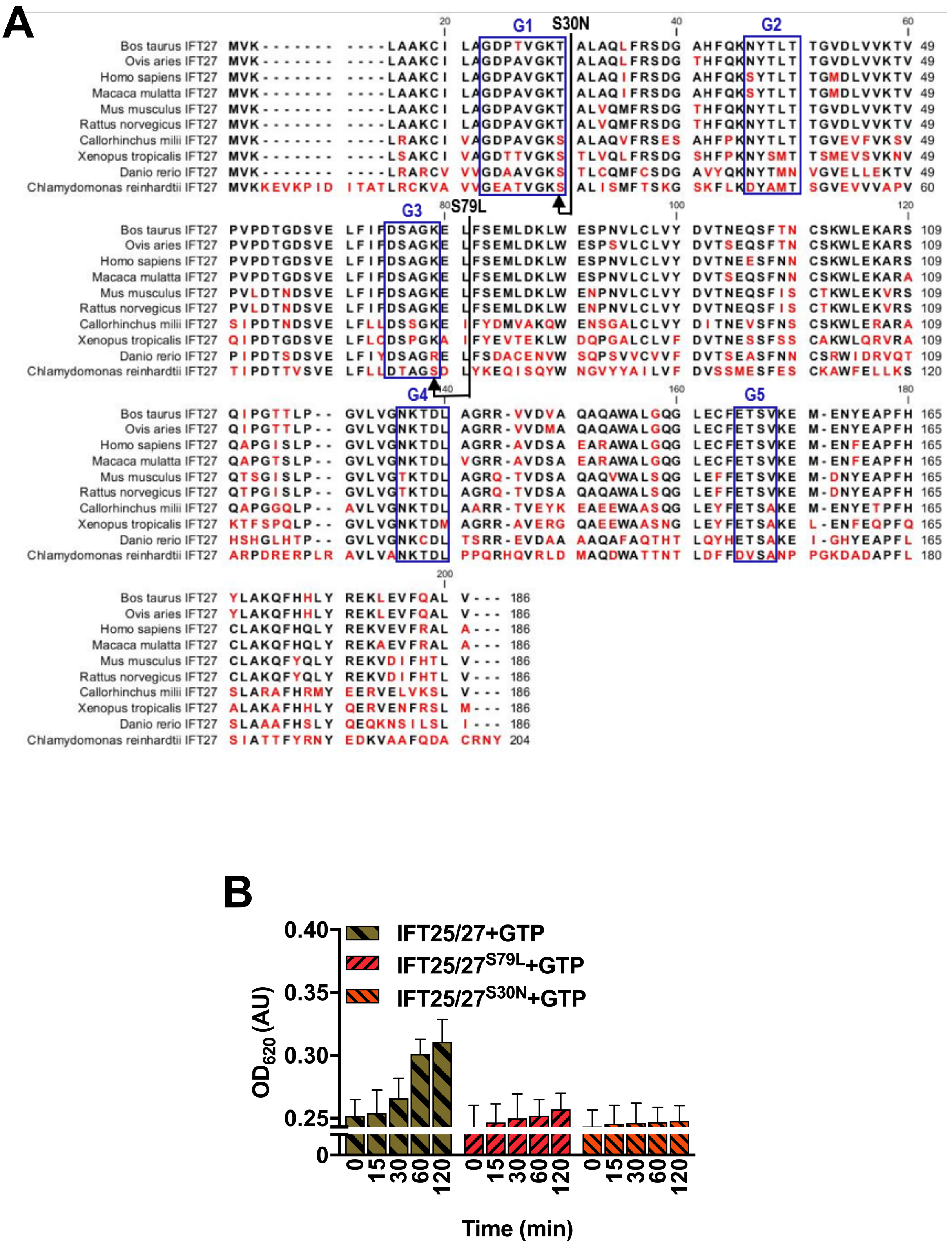
